# Methylthioadenosine is Not Dramatically Elevated in *MTAP*-Homozygous Deleted Primary Glioblastomas

**DOI:** 10.1101/769885

**Authors:** Yasaman Barekatain, Victoria C. Yan, Jeffrey J. Ackroyd, Anton H. Poral, Theresa Tran, Dimitra K. Georgiou, Kenisha Arthur, Yu-Hsi Lin, Nikunj Satani, Elliot S. Ballato, Ana deCarvalho, Roel Verhaak, John de Groot, Jason T. Huse, John M. Asara, Florian L. Muller

## Abstract

**In Brief:** The co-deletion of *MTAP* in the *CDKN2A* locus is a frequent event in diverse cancers including glioblastoma. Recent publications report that significant accumulations of the MTAP substrate, methylthioadenosine (MTA), can sensitize *MTAP*-deleted cancer cells to novel inhibitors of PRMT5 and MAT2A for targeted therapy against tumors with this particular genetic alteration. In this work, using comprehensive metabolomic profiling, we show that MTA is primarily secreted, resulting in exceedingly high levels of extracellular MTA *in vitro*. We further show that primary human glioblastoma tumors minimally accumulate MTA *in vivo*, which is likely explained by the metabolism of MTA by *MTAP*-competent stromal cells. Together, these data challenge whether the metabolic conditions required for therapies to exploit vulnerabilities associated *MTAP* deletions are present in primary human tumors, questioning their translational efficacy in the clinic.

**Highlights:**

- Methylthioadenosine (MTA) is elevated in *MTAP*-deleted cancer cells *in vitro*, which provides a selective vulnerability to PRMT5 and MAT2A inhibitors
- Accumulation of MTA in *MTAP*-deleted cancer cells is predominately extracellular, suggesting active secretion of MTA.
- *MTAP*-deleted primary human glioblastoma tumors show minimal intratumoral elevations of MTA, which is likely explained by secretion and metabolism by *MTAP*-competent stromal cells.

**SUMMARY:** Homozygous deletion of the *CDK2NA* locus frequently results in co-deletion of methylthioadenosine phosphorylase (*MTAP*) in many fatal cancers such as glioblastoma multiforme (GBM), resulting in elevations of the substrate metabolite, methylthioadenosine (MTA). To capitalize on such accumulations, therapeutic targeting of protein arginine methyltransferase 5 (PRMT5) and methionine adenosyl transferase (MAT2A) are ongoing. While extensively corroborated *in vitro*, the clinical efficacy of these strategies ultimately relies on equally significant accumulations of MTA in human tumors. Here, we show that *in vitro* accumulation of MTA is a predominately extracellular phenomenon, indicating secretion of MTA from *MTAP*-deleted cells. In primary human GBMs, we find that MTA levels are not significantly higher in *MTAP*-deleted compared to *MTAP*-intact tumors or normal brain tissue. Together, these findings highlight the metabolic discrepancies between *in vitro* models and primary human tumors and should thus be carefully considered in the development of the precision therapies targeting *MTAP*-homozygous deleted GBM.

## INTRODUCTION

Genomic deletions of tumor suppressor genes are prevalent drivers of tumor progression in myriad cancers but are subverted by their therapeutic inaccessibility via precision oncology. Yet such deletion events often confer less obvious genetic vulnerabilities, which may be therapeutically exploited by identifying their metabolic consequences. Homozygous deletion of *CDKN2A/B* at the 9p21 chromosome (chr9p21) is an early clonal event in tumorigenesis (Beroukhim et al., 2010) that is homogenously distributed in GBM, pancreatic adenocarcinoma, and lung cancer (Hustinx et al., 2005). Within the context of GBM, chr9p21 deletions may exceed 30% of all cases (Parsons et al., 2008). Of particular interest for the development of therapies which capitalize on aberrant tumor metabolism is the collateral deletion of the evolutionarily conserved metabolic enzyme methylthioadenosine phosphorylase (*MTAP*). Due to its proximity to the *CDKN2A* tumor suppressor locus (Kamijo et al., 1997; Serrano et al., 1993), co-deletion of *MTAP* may be observed in 80-90% of all tumors harboring the homozygous deletion of *CDKN2A* (Zhang et al., 1996) (**Figure 1a**). The ubiquity of this event amidst the poor prognosis of cancers such as GBM have thus urged the development of novel therapies that capitalize on downstream vulnerabilities conferred by *MTAP* deletion.

**Figure 1:**
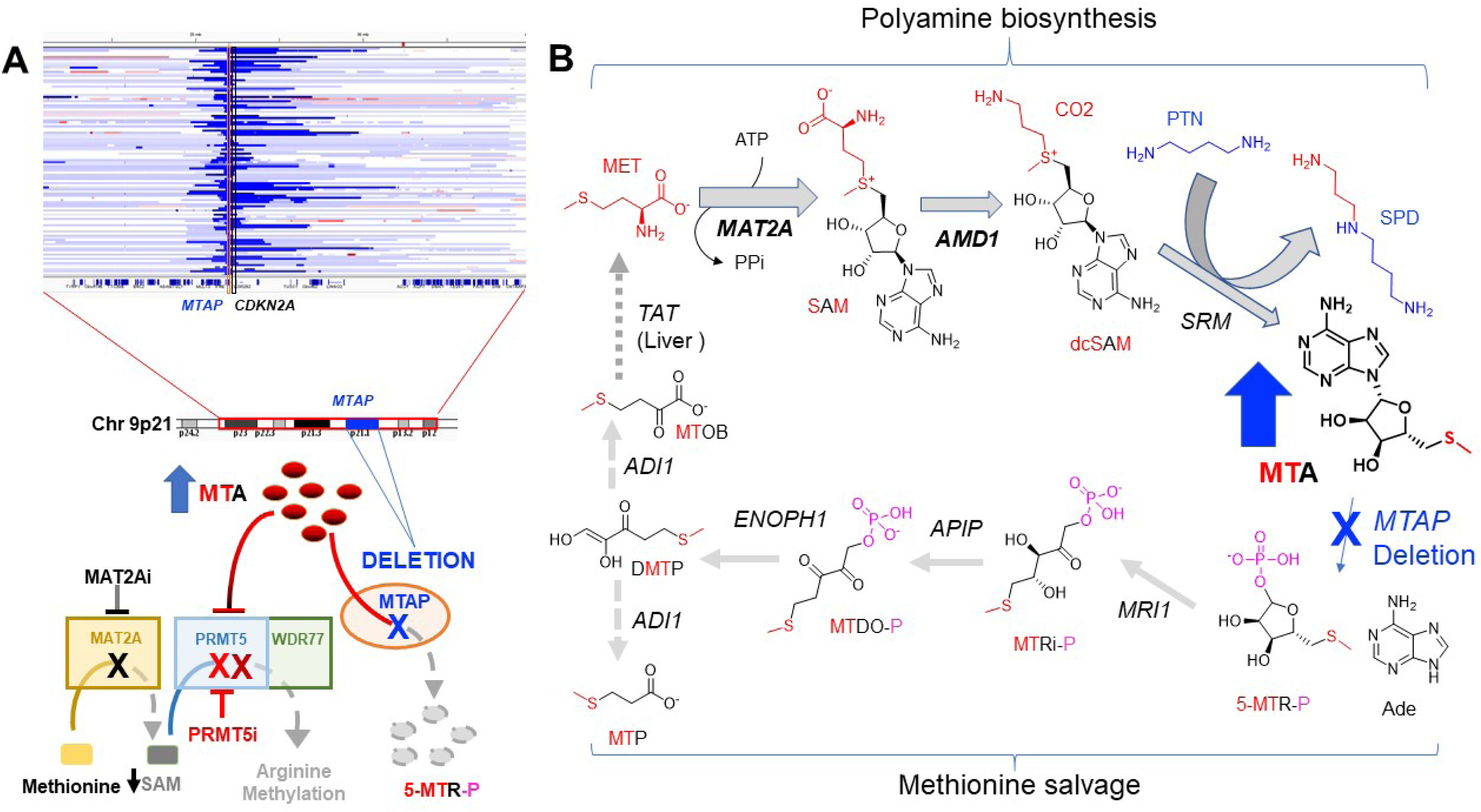
Genomic deletion of *MTAP* as part of the 9p21 locus and the metabolic vulnerabilities associated therewith. **Panel A:** Genomic copy number data (dark blue: bi-allelic or homozygous deletion), from a broad panel of tumors, from the TCGA database, around the tumor suppressor *CDKN2A* in the 9p21 locus. Each strip in the y-axis represents a single tumor, with dark blue showing regions of homozygous deletion, at specific chromosomal location (x-axis). The co-deletion of *MTAP* is expected to result in MTA accumulation. MTA is a structural analogue of SAM (S-adenosylmethionine), and its accumulation can act to sensitize cells with MTAP-deletions to either MAT2A (MAT2A inhibitors, AG270, (Marjon et al., 2016)), inhibitors or PRMT5 (Hörmann et al., 2018; Mavrakis et al., 2016) and even to PRMT1 (Fedoriw et al., 2019; Gao et al., 2019) by acting as a SAM-competitive inhibitor. This competitive inhibition is likely a physiological regulatory feedback mechanism to prevent overproduction of polyamines. **Panel B:** The methionine salvage pathway and the effects of MTAP-deletion. Most genes in the salvage pathway are housekeeping genes that are expressed broadly across cell lines and tissues (DepMap, human protein atlas); TAT and to a lesser extent ADI1, are exclusive liver expressed genes (shown in dashed lines); Pathway drawn based on KEGG. MAT2A is a common essential genes as defined by CRISPR AVANA 19Q3 and is shown in bold (Essential by CRISPR in 600/625 cell lines); ADM1 (CRISPR: 162/625); SRM (CRISPR: 23/625) are cell line selectively essential, MTAP and all downstream genes in the pathway are non-essential according to AVANA 19Q3. Metabolites are color coded to indicate origin of atoms (methionine, red; adenine, black). Abbreviations are as follows. Met: Methionine; SAM: S-adenosylmethionine; dcSAM: S-adenosylmethionineamine; MTA: methylthioadenosine; SPD: spermidine; PTN: putrescine; MTOB: 4-Methylthio-2-oxobutanoate; DMTP:1,2-Dihydroxy-5-(methylthio)pent-1-en-3-one; MTP:3-(Methylthio)propanoate; MTDO-P: 5-(Methylthio)-2,3-dioxopentyl phosphate; MTRi-P: S-Methyl-5-thio-D-ribulose 1-phosphate; 5-MTR-P: Methylthioribose-1-phosphate; Ade: adenosine.

In addition to its broader role in polyamine biosynthesis, MTAP is critically involved in the salvage pathways of both methionine and adenine, catalyzing the conversion of methylthioadenosine (MTA) to S-methyl-5-thio-D-ribose-1-phosphate (**Figure 1b**). Cancers sustaining *MTAP* homozygous deletions are thus expected to accumulate MTA; extensive *in vitro* evidence in diverse cancer cell lines support this logic (Marjon et al., 2016; Mavrakis et al., 2016). Efforts to actionize on this intriguing metabolic phenotype have identified the inhibitory role of excess MTA on protein arginine methyltransferase 5 (PRMT5), a key regulator of transcription. PRMT5 exerts its regulatory effects when complexed with WD repeat-containing protein (WDR77) to generate what is known as the “methylosome” (Friesen et al., 2001, 2002; Karkhanis et al., 2011). High levels of MTA act as an endogenous inhibitor of PRMT5 activity, thereby hindering methylosome formation and rendering cells sensitive to reduced levels of PRMT5 and WDR77.

Therapeutic targeting of PRMT5 in *MTAP*-homozygous deleted cancer cells has thus been upheld as a promising strategy to prevent methylosome organization, selectively killing cancer cells (**Figure 1a**) (Kryukov et al., 2016; Marjon et al., 2016; Mavrakis et al., 2016). Another approach leverages the relationship between PRMT5 and S-adenosylmethionine (SAM) (Chan-Penebre et al., 2015; Marjon et al., 2016; Pathania et al., 2017). SAM is the natural substrate and MTA is a substrate competitive inhibitor of PRMT5. Lowering levels of SAM would thus exacerbate the inhibitory effect of elevated MTA. Accordingly, inhibition of methionine adenosyl transferase (MAT2A), which generates SAM, has also been upheld as a potential therapeutic target in *MTAP*-homozygous deleted cancers (**Figure 1a**) (Chan-Penebre et al., 2015; Marjon et al., 2016; Pathania et al., 2017), which has progressed all the way to an ongoing phase I trial (NCT03435250).

Both of the aforementioned therapeutic approaches are predicated on there being exceedingly high levels of MTA in *MTAP*-deleted cancer cells compared to *MTAP*-intact tissues. While it may seem natural that *MTAP*-homozygous deleted primary human tumors should also display this phenotype, recent reports identifying this intriguing vulnerability have not actually measured the levels of MTA in primary human tumors (Marjon et al., 2016; Mavrakis et al., 2016). In fact, we could not find any studies in the literature that reported MTA measurements in actual primary human tumors with an established *MTAP* genotype. Through combined analysis of both public domain data from previous metabolomics profiling studies and our own data generated in *MTAP*-homozygous deleted primary human GBMs, we found that the dramatically elevated levels of MTA found in glioma cell lines cannot be extrapolated to primary GBMs. A series of metabolomics studies in cell culture and on primary tumors converge on the observation that accumulation of MTA in culture is a predominately extracellular phenomenon, suggesting secretion out of *MTAP*-deleted cells. Expounding on this finding in primary tumors, our data strongly suggest that the abundance of non-malignant *MTAP*-WT stromal cells metabolize the secreted MTA from the *MTAP*-homozygous deleted GBM cells. As the promise of synthetic lethal therapies against *MTAP*-homozygous deleted cancers are premised on vast, intracellular accumulations of MTA, our data caution against the expedient translation of the *MTAP*-deletion targeted precision therapies to the clinic and strongly contend for more research into the fundamental metabolic differences between model systems and primary tumors.

## RESULTS

### *In vitro* MTA accumulation in *MTAP*-deleted cells is primarily an extracellular phenomenon

MTA is a small, neutral, sulfur-containing nucleoside, which suggests that it may permeate the cell membrane through a nucleoside transporter or perhaps even via passive diffusion. Metabolic profiling studies using independent metabolomics platforms have uniformly concluded that intracellular levels of MTA are elevated >20-fold *in vitro* (Batova et al., 1996; Iizasa et al., 1984; Kamatani and Carson, 1980; Li et al., 2019; Marjon et al., 2016; Stevens et al., 2009, 2010). However, the extent to which these studies clarify between intracellular and extracellular distribution is not always apparent. In the studies that have differentiated between the two, increases in MTA in *MTAP*-homozygous deleted versus WT cell lines are actually quite modest (Ortmayr et al., 2019). For instance, in the NCI-60 profiling study of Ortmayr *et al*, which focused exclusively on intracellular metabolites (extensive washing of cell pellet), there is only ~15% increase in MTA in *MTAP*-deleted compared to WT cancer cell lines (**Supplementary Figure S1a**). However, a collaboration between Metabolon Inc and the NCI using the NCI-60 panel did not specify whether washing of the cell pellet was performed and reported a 3-fold increase in MTA in *MTAP*-deleted compared to intact cell lines (**Supplementary Figure S1b**). Even more strikingly, re-analysis of one notable study that conducted mass-spectrometry profiling on conditioned media (secreted metabolites); showed >100-fold increases in MTA in *MTAP*-deleted cell lines compared to intact cell lines (**Supplementary Figure S1c**; for comparison with lactate, see **Supplementary Figure S1d** (Dettmer et al., 2013)). The micromolar extracellular levels of MTA reported sharply contrasts the nanomolar to picomolar levels of MTA typically found in conditioned media of *MTAP*-WT cells (Kirovski et al., 2011; Stevens et al., 2010). Suffice to say, micromolar concentrations of extracellular MTA in are biochemically outstanding. Indeed, we find that MTA is abundant enough in conditioned media of *MTAP*-deleted U87 glioma cells, to be detectable by relatively insensitive techniques such as NMR (see below). Though the general trend of increased levels of MTA in *MTAP*-deleted cell lines was consistent across all experiments, the magnitude of intracellular accumulation was strongly dependent on how extensively the cells were washed prior to metabolite extraction. Less vigorous washing of the cell pellet may give the appearance of high levels of intracellular MTA by inadvertently including the abundant extracellular MTA in the media.

To independently investigate the likelihood of export from *MTAP*-deleted glioma cells, we compared the concentrations of MTA from the intra- and extracellular environments of three verified *MTAP*-null cell lines: U87, Gli56, SW1088 (**Figure 2a**). Both the extensively washed cell pellet and conditioned media were profiled using the Beth Israel Deaconess Medical Center (BIDMC) platform. Strikingly, the modest intracellular increases of MTA, were vastly overshadowed by extracellular levels MTA, again evidencing secretion (**Figure 2b vs. 2c**). To corroborate our findings, we also performed a time-course experiment for *MTAP*-deleted and *MTAP*-WT cell lines. At given time points, we added new media before profiling both the intracellular (cell pellet) and extracellular metabolites (conditioned media). Though a time-dependent increase was apparent in both the metabolite profiles, it was exacerbated in the extracellular profile (**Figure 2d**). This imbalance further supported active MTA secretion. A comparison of intracellular and extracellular levels of MTA, adenosine (nucleoside precursor), SAM and the structural analogue of MTA, methyladenosine, clearly demonstrates the tight regulation of intracellular retained versus secrete metabolites (**Figure 2d-g**). Unlike the exclusively intracellular presence of adenosine and SAM, MTA and methyladenosine gradually accumulate extracellularly, suggesting that both are actively extruded.

**Figure 2:**
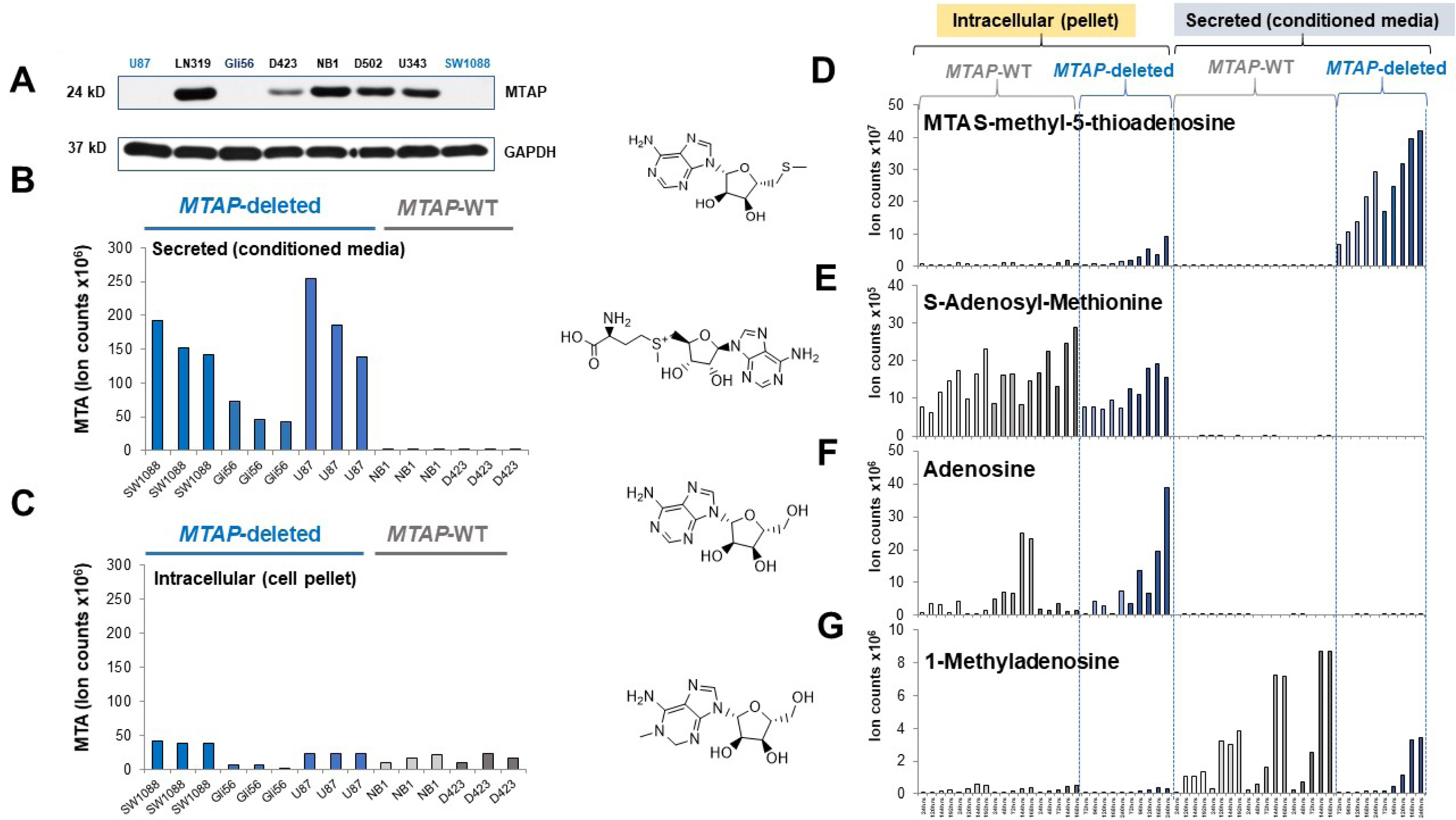
Imbalance of MTA between intracellular and extracellular levels of MTA in MTAP homozygously deleted cells. All metabolite measurements shown were obtained using the BIDMC platform. Levels of MTA in a panel of glioma cell lines differing in MTAP-deletion status (confirmed by western blot, **Panel A**) for both conditioned media (**Panel B**) and washed pellet (**Panel C**). Each bar represents one biological replicate. **D-G**: Both WT and *MTAP* homozygous deleted glioma cells were grown for the indicated time points and harvested for polar metabolites. To prevent media contamination, cells on plates were washed 3x with cold PBS, then spun down at 1,500 g, and extracted with methanol. Each cell line thus has a paired sample of intracellular (pellet) and extracellular metabolites (conditioned media). The amount of media harvested was adjusted so that the total ion count (total metabolites) from both sets of samples would be the same; color coding reflects the same cell line for pellet/media. Intracellular and extracellular levels of MTA for each profiled cell line are aligned with S-adenosylmethionine, adenosine and 1-methyladenosine for comparison. X-axis: time of harvest after last media change; Y-axis, ion counts for each metabolite. Each bar represents one biological replicate. There is a distinct, time-dependent increase in MTA in media in *MTAP*-deleted cells. Though intracellular levels of MTA also increase over time, this elevation is much more modest. Indeed, there is a striking imbalance of MTA in the pellet vs the media; this is in contrast to what is observed with Adenosine and S-adenosyl methionine which are exclusively intracellular.

While the precise mechanism of MTA transport remains underexplored, its identity as an uncharged, sulfur-containing nucleoside makes passive diffusion or movement by adenosine transporters likely (Iizasa et al., 1984; Munshi et al., 1988; Plagemann and Wohlhueter, 1980). That adenine transporters are known to have broad substrate specificity further substantiates this explanation (Munshi et al., 1988; Plagemann and Wohlhueter, 1980). For comparison, the anionic metabolite, 2-hydroxyglutarate, has a distinctly intracellular presence, owing to its highly polar nature and absence of any substrate-specific transporters to facilitate its exit out of the cell. Both our data and structure-based-logic are supported by similar observations in other cancer models (Kamatani and Carson, 1980; Kirovski et al., 2011; Limm et al., 2014; Stevens et al., 2010). In fact, direct measurement of MTA secretion into culture medium out of *MTAP*-deleted cells was found to occur at a rate of 0.58 to 0.70 nmol/hr/mg of protein in a leukemia cell culture model (Kamatani and Carson, 1980) compared to < 0.1 for *MTAP* WT leukemia. Paired with these previous studies, our findings build a compelling case for MTA export in *MTAP*-deleted glioma cells and likely all cancer cell lines.

### Metabolomic profiling data from independent platforms do not corroborate a strong elevation of MTA in *MTAP*-homozygous deleted primary human GBM

Metabolomics analyses of primary human GBM tumors are sparsely reported in the literature. The few studies that have been conducted have favored comparison between tumor populations at different stages or grades, rather than considering the unique metabolic landscape of individual tumors (Erb et al., 2008; Hakimi et al., 2016; Prabhu et al., 2019; Sahu et al., 2017; Xu et al., 2019). In accordance with the proclivities of the field, we first queried supplemental data from studies using the Metabolon Inc platform on primary human GBMs (Chinnaiyan et al., 2012; Prabhu et al., 2019). As these studies were focused on identifying differences in metabolites between different glioma grades, no genomics of individual tumors were available. We thus sought to match the metabolomics data on different glioma grades with the reported frequency of *MTAP* deletions in different glioma grades. Previous studies have shown that the incidence of *CDK2NA*/*MTAP* homozygous deletion in grade II glioma is less than 3%; for grade II/III glioma it is never more than 10%. In sharp contrast, the frequency high as 50 percent in GBM (Brennan et al., 2013; Cancer Genome Atlas Research Network et al., 2015; Parsons et al., 2008). We would thus expect that the matched elevations of MTA would correspond to the aforementioned deletion frequencies per tumor grade, with higher levels of MTA in GBM and considerably lower levels especially in grade II glioma. However, despite these discordant frequencies in *MTAP*-homozygous deletion status, analysis of MTA levels from among different glioma grades actually shows similar levels of MTA in grade II/III, grade III and grade IV (GBM) tumors (Chinnaiyan et al., 2012) (**Supplementary Figure S3a,b**). Importantly, no extreme outliers in MTA were evident. At best, the highest levels of MTA in individual GBM cases are just about 2-fold higher than the median of (MTAP-WT) lower grade glioma tumors. This evidently runs contrary to the expected phenotype in *MTAP*-homozygous deleted tumors. For comparison, these data also run in stark contrast to the extreme elevations (>100-fold) in 2-hydroxyglutarate driven by IDH mutations (**Supplementary Figure S3c**). To corroborate these findings, we analyzed metabolomics data obtained from the Metabolon Inc platform (Prabhu et al., 2018), which neither showed extreme outliers in GBM-associated MTA nor higher levels of MTA comparing lower grade glioma (**Supplementary Figure S4a**).

To extend our *in vitro* observations on secretion to the *in vivo* environment, we sought to identify any notable change in MTA levels in cerebrospinal fluid (CSF) of GBM versus lower grade glioma or normal brain. No ^1^H NMR based metabolomic study that we could find actually reported increases of MTA in GBM (Shao et al., 2014). Surprisingly, a study using the BIDMC platform only reported marginal differences between MTA levels in the CSF collected from GBM and that collected from the normal brain (Locasale et al., 2012) (**Supplementary Figure S4b**). Given that the frequency of *MTAP* homozygous deletions in GBM is at least 35%, we would have expected to observe a dramatically higher level of MTA in GBM CSF compared to normal brain CSF. Although there was a 1.5-fold difference between GBM and normal brain, with a p-value of 0.0407 by t-test (Table 2 in (Locasale et al., 2012)), this difference becomes non-significant when correcting for multiple hypothesis testing. Even if these data do suggest an underlying trend, this is a far cry from the greater-than-100-fold elevation in secreted MTA for *MTAP*-deleted cells in culture (**Figure 2, Supplementary Figure S1**).

To answer the question directly, we analyzed MTA levels from our metabolomic profiling data generated with the BIDMC platform in primary tumors with full genomic annotation (Wang et al., 2017) for direct comparison between GBM tumors of verified *MTAP*-deleted and *MTAP*-WT genotype. This is the first study that directly addressed the effects of *MTAP*-deletion in primary human tumors. We find that there is a marginal difference (~1.7-fold higher median levels of MTA when considering the raw data, and ~1.3-fold higher when corrected for loading; **Figure 3a, b**) between the average levels of MTA in *MTAP*-intact and *MTAP*-deleted tumors. Though the individual differences in levels of MTA among the tumors within each population appear to overshadow the seemingly small differences between the two populations, the overall difference between the two groups did meet statistical significance (p < 0.03; by unpaired 2-tailed t-test with unequal variance). However, we note that this marginal elevation pales in comparison to outliers driven by specific genetic events that fully recapitulate metabolic data recorded intracellularly in cell culture, such as the >100-fold increase in the levels of 6-phosphogluconate and 2-hydroxyglutarate, driven by the homozygous deletion of *PGD* (Molina et al., 2018; Sun et al., 2019) and the point mutation of IDH1, respectively (**Figure 3c,d; and Supplementary Figure S3c, S4c**).

**Figure 3:**
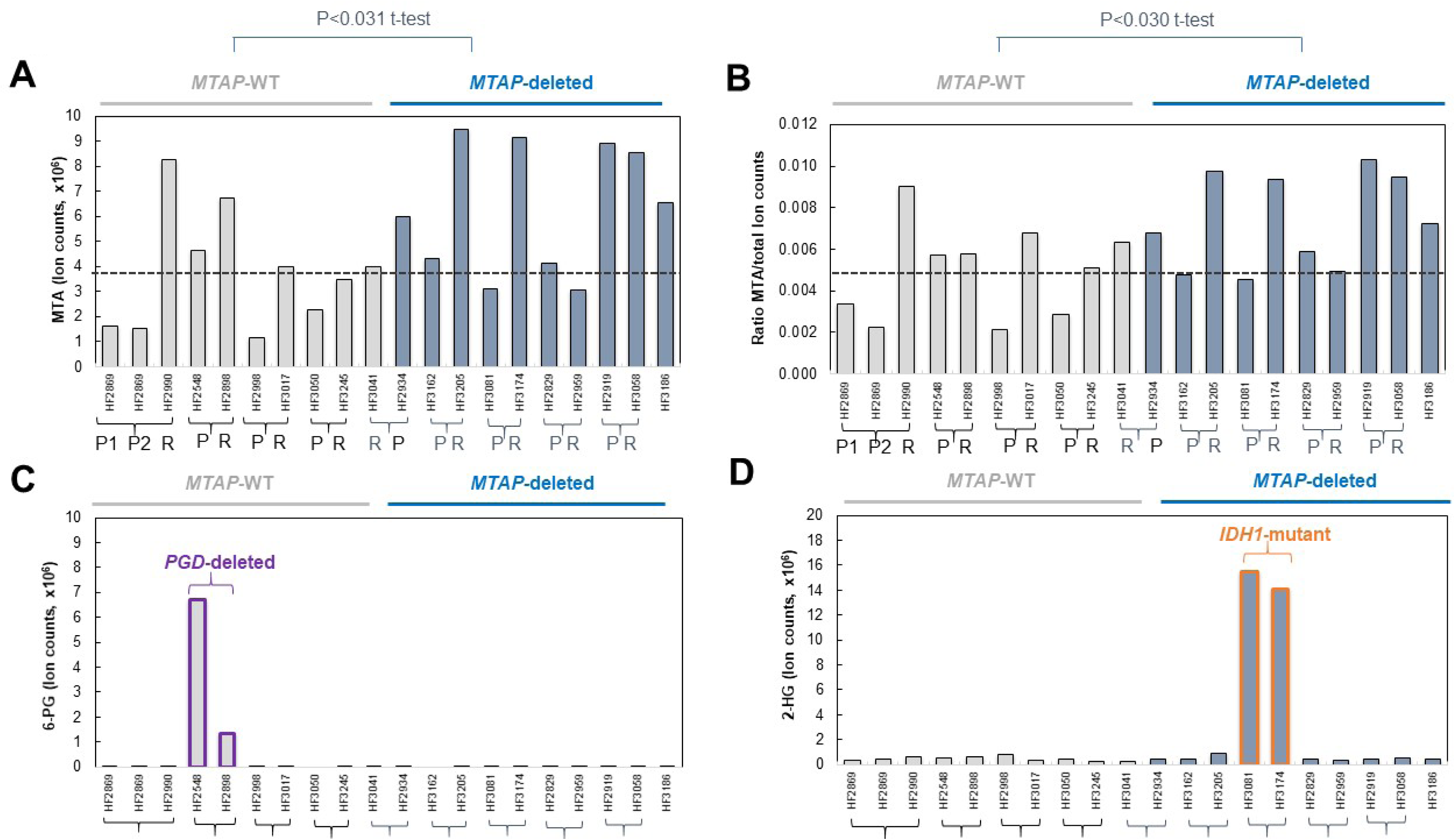
Minimal elevation of MTA in *MTAP*-deleted human GBM tumors. **Panel A**: MTA levels (absolute ion counts) in *MTAP*-deleted versus *MTAP*-intact primary (P) and recurrent (R) GBM tumors from the same case (bracket), using the BIDMC metabolomic platform, that were MTAP-intact (grey) and MTAP-deleted (blue). The median MTA of MTAP-intact tumors is shown as a grey dashed line. **Panel B**: Same data but expressed as a ratio of the total ion count of each tumor, for sample loading normalization. P values unpaired t-test, with un-equal variance, indicated. **Panel C**: Metabolite levels of 6-phosphogluconate (6-PG) derived from the same profile; tumors with homozygous deletion of 1p36, which includes 6-phosphogluconate dehydrogenase (*PGD*) is indicated. **Panel D**: 2-hydroglutarate (2-HG) show dramatic elevation in the paired cases with IDH1-mutant tumors. Genomic data for this tumor series were previously published (Wang et al., 2017).

In addition to these mass spec metabolomics studies, large scale (n >100 cases) NMR, HR-MAS metabolomic profiles of GBM have not detected methylthioadenosine in *MTAP*-deleted tumors even though HR-MAS can usually detect metabolites at a minimum concentration of 1 µM (Elkhaled et al., 2014). That MTA could still not be detected in these contexts suggests that that less than 1 µM of MTA is present in GBM tumors, which concurs with our conclusion that there are no tumors, regardless of *MTAP*-deletion status, have outliers in MTA.

### Primary GBM tumors are extensively populated by MTAP-expressing stromal cells

To both verify the diagnosis of *MTAP*-homozygous deletion and contrast this with the extensive levels of MTAP expression in non-malignant tissue, we performed immunohistochemistry. It is already well-established that GBM tumors may have up to 70% stromal cells, including non-transformed reactive astrocytes, lymphocytes, endothelial cells, fibroblasts, and axonal and neuronal remnants (Kim et al., 2015; Wang et al., 2017). We first validated an MTAP rabbit monoclonal antibody (Clone: [EPR6892], ab126623)) by demonstrating strong staining in formalin fixed paraffin embedded (FFPE) sections of *MTAP*-WT and loss-of-staining in *MTAP*-deleted xenografted tumors established in known *MTAP*-genotype cell lines (**Supplementary Figure S5–S7**). Every xenograft established from *MTAP*-WT (or isogenic MTAP-rescued) cancer cell lines showed strong staining while all xenografts established from MTAP-deleted cell lines showed no staining (**Supplementary Figure S5–S7**). At the same time, non-tumor stromal cells such as microglia, lymphocytes, and endothelial cells stained positive for MTAP. Together, these staining results support that this antibody accurately quantifies MTAP levels in FFPE slides. Notable among our stained slides was the significant amount of positive staining in normal mouse brain and stromal cells, especially given that xenografts typically grow much less invasively and are less populated by stromal cells compared to human GBMs. By extension, true human GBMs are thus more likely to contain a higher proportion of stromal components such as astrocytes and microglia.

Next, we applied this antibody to primary GBM tumor FFPE sections. Of 60 cases analyzed, we find that approximately 40% of cases show complete lack of MTAP staining in GBM tumor cells, which corresponds to the expected MTAP-deleted frequency in GBM, (Brennan et al., 2013); notably, strong staining was observed in non-malignant stromal cells. The remaining cases show strong, uniform staining in both glioma and stromal cells (representative stained cases are shown in **Figure 4**, **Supplementary Figure S8**). The histology of the MTAP-negative primary GBM cases looks quite similar to the *MTAP*-deleted intracranial xenografted cases, with the exception that the degree of *MTAP*-positive stromal infiltration is much greater in the primary human GBM tumors compared to xenografts (**Supplementary Figure S8** provides a sampling of Primary GBM cases differing in the degree of stromal content). The lower *MTAP*-positive stromal content in xenografts is of high relevance as this indicates that xenografted tumors likely underestimate the problem of stromal cell dependent metabolism of tumor cell secreted MTA (**Figure 4c**).

**Figure 4:**
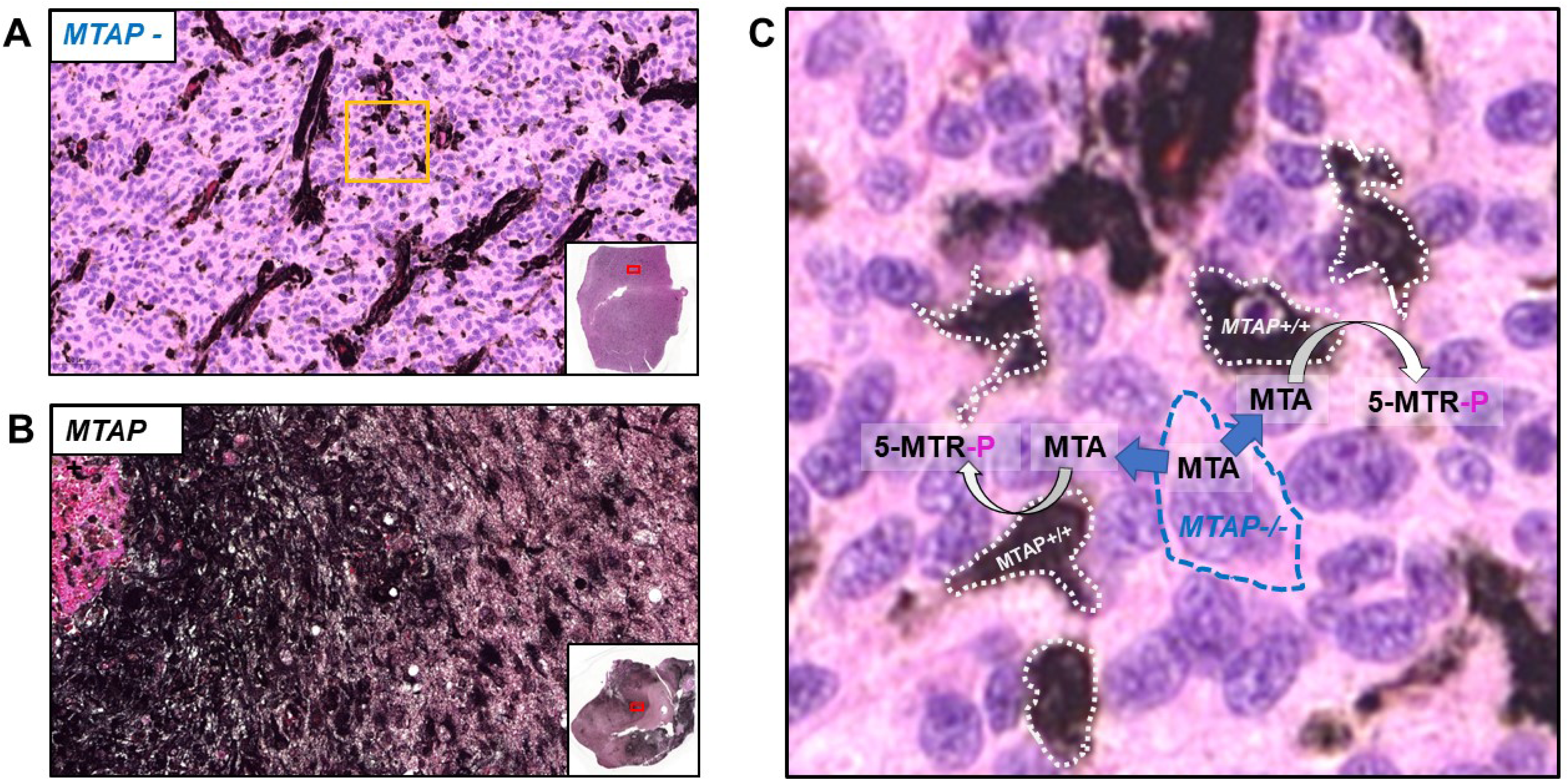
Human GBM tumors are extensively infiltrated by MTAP-expressing non-malignant stromal cells. IHC was performed with rabbit monoclonal anti-MTAP (ab126623; EPR6892) and the slides were developed with EnzMet (black, MTAP expression) and counterstained with hemoatoxylin (blue, nuclei) and eosin (pink, cytosol and extracellular space). Representative sections of MTAP-negative (**A**) and an MTAP-positive GBM tumor (**B**); 20X objective. Note the extensive MTAP-positive staining areas on the background of otherwise non-staining MTAP-negative glioma cells; histopathological evaluation of morphology indicates that these MTAP-positive cells correspond to stromal components, including microglia, endothelial cells and activated astrocytes. **Panel C**: Higher magnification view of Panel A golden square. Any MTA secreted by *MTAP*-deleted glioma cells (blue dashed outline), stands to be phosphorylated by MTAP (producing 5-MTR-P) either released or taken up by MTAP-expressing stromal cells (white dashed outlines).

### Co-culture of *MTAP*-deleted with *MTAP*-rescued glioma cell lines abrogates MTA accumulation in conditioned media

We then co-cultured U87 (*MTAP*-deleted) with U87 MTAP-isogenic rescued cells and determined the effects of MTA accumulation in the media to better model the situation within *MTAP*-deleted tumors (**Figure 4**). Using ^1^H-NMR, we measured robust accumulation of MTA in the conditioned media of U87 cells but not D423 (MTAP-intact), or U87 pCMV MTAP (*MTAP*-rescued glioma) cells (**Supplementary Figure S9**). When U87 parental *MTAP*-deleted glioma cells were co-cultured with U87 MTAP-rescued cells, no MTA is detectable by NMR in conditioned media (**Supplementary Figure S10**). This indicates that either U87 MTAP-rescued cells release functional MTAP enzyme into the media or they take up MTA from the media and metabolize it. Either way, this confirms that MTAP-intact cells in the vicinity of *MTAP*-deleted cancer cells deplete the extracellular accumulation of MTA, negating the selective vulnerability to PRMT5/MAT2A inhibitors conferred by MTAP deletion.

## DISCUSSION

We initially embarked on this study in response to the exigence of *MTAP*-deleted precision therapy endeavors despite lack of exploration of the metabolic state of primary tumors. As the efficacy of *MTAP*-deletion directed therapies hinges on massive intracellular accumulation of MTA (Fedoriw et al., 2019; Gao et al., 2019; Hörmann et al., 2018; Kryukov et al., 2016; Marjon et al., 2016; Mavrakis et al., 2016), ensuring that this *in vitro* phenotype is consistent in human tumors seemed to be a natural and urgent area for investigation. Perhaps surprisingly, none of the recent papers documenting MTAP-dependent vulnerabilities actually reported measurements of MTA in primary human tumors or even xenographed tumors of defined *MTAP*-deletion status, despite the significant efforts that have been built on the supposition that observed elevations in MTA *in vitro* would be reflected in primary tumors (Fedoriw et al., 2019; Gao et al., 2019; Hörmann et al., 2018; Kryukov et al., 2016; Marjon et al., 2016; Mavrakis et al., 2016).

We took to the literature—keen on finding studies describing MTA levels in distinct, genetically defined *MTAP*-deleted primary tumors—but with little success. *<u>We could not find any reports in the literature directly measuring levels of MTA in genomically-identified MTAP-deleted versus MTAP-intact primary human tumors, whether GBM or others.</u>* This was quite surprising, given that MTA is a metabolite frequently detected in many metabolomic profiling studies. We queried the public domain metabolomic datasets that have been generated using multiple different platforms (Chinnaiyan et al., 2012; Locasale et al., 2012; Sanderson et al., 2019). While we were unable to locate metabolomic datasets with companion genomic information, we found multiple high quality metabolomic datasets for primary glioma of different grades. Given the high prevalence of *MTAP* homozygous deletions in grade IV-glioma/GBM (~40%) and contrastingly low frequency in lower grade glioma, comparison of these two tumor sets can proves insightful. If *MTAP* deletions do indeed lead to MTA accumulation in primary human tumors, we would expect to observe dramatically higher levels of MTA (>20-fold in data from intracellular data *in vitro* from (Marjon et al., 2016) Figure 3C in that publication) in a substantial fraction of GBM tumors. *<u>Data from three independent metabolomic datasets generated from different platforms indicate that the differences in MTA levels, even when considering outliers, are at best, 2-fold higher in purportedly MTAP-deleted versus MTAP-intact tumors.</u>* We then interrogated one of our own metabolomic datasets generated from primary and recurrent snap frozen GBM tumors utilizing the BIDMC platform with complementary genomic data (Wang et al., 2017). Even if our data may allow for the possibility of marginal difference in levels of MTA between *MTAP*-deleted and *MTAP*-WT tumors (1.7-fold or 1.3-fold increase, depending on normalization; p < 0.03), there is extensive overlap between the groups and this is nowhere near the vast elevations in MTA reported for glioma cell lines *in vitro*.

In the literature, we noticed strong discrepancies in the magnitudes of reported MTA-increase in *MTAP*-deleted versus *MTAP*-intact cancer cell lines from different studies (**Supplementary Figure 1**). Careful analysis of these results in light of our own *in vitro* data evidencing far greater accumulations of extracellular, rather than intracellular, MTA points to MTA secretion out of *MTAP*-deleted cells and the discrepancies from different studies arising from how well intracellular versus extracellular metabolite levels are separated during sample processing. Indeed, similar observations have been previously reported (Kamatani and Carson, 1980; Riscoe and Ferro, 1984), which further corroborate this finding. Another study also observed extracellular accumulation of MTA in *MTAP*-deficient cells, but only when cultured with heat-inactivated fetal bovine serum (FBS) or fetal horse serum, which does not contain active MTAP (Riscoe and Ferro, 1984). Heat-inactivated FBS is currently the standard for cell culture and was thus used in our studies and all other recent MTAP-related studies.

In sum, our findings on massive extracellular accumulations do not extend to our metabolome analyses of *MTAP*-deleted primary GBM tumors or CSF: at best, marginal elevations in MTA were observed in any of these. A variety of reasons may explain the discrepancy between cell culture and human tumor data. The simplest possible explanation is that the exceedingly high levels of reported intracellular MTA may be due to less vigorous washing of the cell pellet prior to metabolomic profiling. Biochemical differences between cell culture models and the *in vivo* environment may also be a contributor: MTAP is present in both human plasma (Carrera et al., 1986; Wishart et al., 2007) and in the CSF (Guldbrandsen et al., 2014), making the conversion of released MTA to S-methyl-5-thio-D-ribose-1-phosphate and adenine possible. Primary tumor environments are also characterized by high stromal content comprised of *MTAP*-WT cells (**Figure 4**). As MTA is a neutral nucleotide, both passive diffusion and movement via adenine transporters (Limm et al., 2014; Munshi et al., 1988) into *MTAP*-WT cells are possible, though the latter is much more likely (Iizasa et al., 1984; Munshi et al., 1988; Plagemann and Wohlhueter, 1980). Under this explanation, the net movement and metabolism of MTA by stromal cells makes the absence of any detectable increase in MTA in primary GBMs and CSF of MTAP-deleted tumors self-evident. Finally, we also considered the possibility of discordant levels of the enzymes preceding the MTAP reaction—adenosylmethionine decarboxylase and spermidine or spermine synthase— between cultured cells and primary GBMs. However, a survey of the Human Protein Atlas shows that expression is comparable amongst the two (e.g. Human Protein Atlas: SRM).

A combination of the above explanations most likely contributes to the cultured cells versus primary tumor discrepancy. It is already well-established that cell culture conditions do not always accurately reflect physiological conditions (Ackermann and Tardito, 2019; Sanderson et al., 2019; Vande Voorde et al., 2019). Prior to *in vitro* metabolomics profiling, we cultured our cells using heat-inactivated fetal bovine serum which, thereby nullifying any *MTAP*-activity that would otherwise be present in the serum. The more significant contributor to the primary GBM results may be attributed to the abundance of stromal cells, which may equal or surpass the number of malignant cells— even in very aggressive tumors (**Figure 4**). While the metabolomic data are strictly limited to GBM, the secretion versus intracellular accumulation of MTA is evident in cell lines of other cancer types (**Supplementary Figure S1**) (Su et al., 2011), and the presence of MTAP in extracellular biological fluids has been documented in whole blood (Ruefli-Brasse et al., 2011).

Ultimately, these findings are significant because precision therapies targeting *MTAP*-deleted tumors are predicated on high levels of MTA for efficacy. Reports of high extracellular levels of MTA in *MTAP*-deleted cells permeate the early biochemical literature. Our results corroborate and, with the advent of metabolomics, offer a more meticulous inspection of global metabolite changes across a range of cell lines and primary GBM tumors. That *MTAP* deletions can be leveraged as a point of selective vulnerability in myriad cancers has spurred much excitement into finding targetable proteins in its metabolic network. Accordingly, MAT2A has emerged as the pre-eminent target thus far (Kryukov et al., 2016; Mavrakis et al., 2016). In addition to our aforementioned shortcomings, targeting MAT2A raises a number of independent issues. MAT2A catalyzes the production of SAM and SAM is evidently essential for biological function on its own. CRISPR/Avana datasets show that, MAT2A (and PRMT5, for that matter) is an essential gene, irrespective of *MTAP* deletion status (summarized in **Figure 1**).

Several other strategies that do not require accumulation of MTA have been proposed to exploit the *MTAP*-homozygous deletion for the purposes of precision oncology. Specifically, the purine biosynthesis of methionine biosynthesis defects (Kindler et al., 2009; Ruefli-Brasse et al., 2011; Tang et al., 2018). But again, the issue of *MTAP*-WT-expressing stromal cells or MTAP present in extracellular biological fluids is also problematic for *in vivo* efficacy for these alternative strategies. These hurdles beg the question as to whether the high frequency of *MTAP* deletions can even be therapeutically actionable in the clinic.

To this end, we offer three different strategies that might overcome the issue of MTA secretion. First, the most obvious is to identify the nucleoside transporters and generate appropriate inhibitors. The second possible solution involves inhibition of extracellular MTAP, if this proves to be as significant a route of MTA metabolism as other have suggested (Ruefli-Brasse et al., 2011). Though the latter may appear to be a lofty aspiration, we contend there are two fairly straightforward approaches to accomplishing this. First, as the MTAP reaction is a phosphorolysis, it is reasonable to conclude that an MTAP phosphonate transition state analogue may effectively inhibit extracellular MTAP. Transition state inhibitors have, indeed, already been synthesized (Singh et al., 2004). However, the absence of an anionic phosphate or phosphonate moiety makes these inhibitors cell permeable. This negates the exclusive extracellular MTAP inhibition that would have otherwise occurred had a phosphate or phosphonate moiety been present in these structures. Phosphonates and phosphates are poorly cell permeable (Leonard et al., 2016; Lin et al., 2018), but distribute well in water, making them ideal to incorporate in inhibitors of extracellular enzymes. A second approach to selectively inhibiting extracellular MTAP would be a neutralizing monoclonal antibody. Such antibodies have been successfully designed for many growth factors and cytokines, and even enzymes (Farady et al., 2007). Given the near-exclusive extracellular activity of therapeutic antibodies, this may be a viable strategy to neutralize extracellular MTAP-activity and allow MTA to accumulate in *MTAP*-deleted tumors, thus enabling selective sensitization to inhibitors of PRMT5 and MAT2A. Finally, there may be metabolic methods to force overproduction of MTA. For example, *in vitro* studies indicate that supplementation of cells with putrescine can increase MTA secretion rates (Kamatani and Carson, 1980). This is because putrescine is the substrate of spermine synthase, which generates both spermine and MTA (**Figure 1b**). Because putrescine is minimally toxic (LD_50_, > 500 mg/kg; TOXNET CASRN: 110-60-1), it could potentially be used pharmacologically for the deliberate exaggeration of MTA accumulation in *MTAP*-deleted tumors.

## STAR METHODS

### Identification of cell lines with *MTAP*-homozygous deletions

Scoring of homozygous deletions even with robust copy number data can be problematic due to uneven ploidy across cell lines, as well as where deletion boundaries only partially cover the gene, yet eliminate its functional expression. Thus, we sought to identify MTAP functionally null cell lines. For the NCI-60 cell line comparisons, we gathered data from the Sanger Center COSMIC; this data is corrected for ploidy by the ASCAT algorithm (Van Loo et al., 2012). We utilized confirmed data using cBioPortal with the NCI-60 panel data (Cerami et al., 2012; Reinhold et al., 2012), and DepMap CCLE as well as the Genentech collection (Klijn et al., 2015). Where deletion calls were ambiguous (e.g. SF-295), we took at mRNA expression data or literature western blot data (Hörmann et al., 2018; Kryukov et al., 2016; Marjon et al., 2016; Mavrakis et al., 2016) to decide what constituted a functionally MTAP-null cell line. In the NCI-60 cell line panel, the following cell lines lack *MTAP*: NCI-H322, SK-MEL-5, K-562, SF-268, MALME-3M, ACHN, MDA-MB-231, OVCAR-5, SF-268, CCRF-CEM, SR, MCF-7, and A549.

### Cell Culture and Xenograft Generation

The cell lines used in this study that are MTAP-WT: D423 (H423/D423-MG, kindly provided by D. Bigner (Duncan et al., 2010), D502 (H502 in (Duncan et al., 2010)), U343 (U343-MG in (Nistér et al., 1987)), NB1, LN319 (a sub-clone of LN-992 in (Bady et al., 2012)) and the *MTAP*-deleted cell lines are U87, Gli56 (D. Louis), SK-MEL-5 and SW1088. The cell lines SW1088, U87, U343, LN319, SK-MEL-5, NB1 were obtained from the Department of Genomic Medicine/IACS Cell Bank. All cell culture experiments were conducted according to provider’s instructions and as previously described (Leonard et al., 2016). The U87 *MTAP*-rescued cell line was generated utilizing the same procedure we previously utilized to generate an *ENO1*-rescued cell line (PMID: 22895339). The MTAP cDNA was obtained from Lifetechnology and sequenced verified and cloned into pCMV GFP lentiviral vector using Gateway cloning technology. 293T cells were transfected with viral plasmids using polyethylenimine (PEI). The viral supernatant was collected after 72 hr after transfection and added to the U87 cell line for infection.

All cells were cultured at 37° C, in 5% CO_2_ air atmosphere and pH 7.4, in ATCC suggested media (DMEM) unless stated otherwise. DMEM Media has 4.5 g/L glucose, pyruvate and glutamine (Cellgro/Corning #10-013-CV) with 10% (20% for Gli56), fetal bovine serum (Gibco/Life Technologies #16140-071) and 1% Pen Strep (Gibco/Life Technologies#15140-122) and 0.1% Amphotericin B (Gibco/Life Technologies#15290-018). All cell lines were confirmed as myco-plasma negative by ELISA using the MycoAlert PLUS detection kit (Lonza) and were authenticated by short tandem repeat DNA fingerprinting and chromosomal analysis by the Characterized Cell Line Facility and the Molecular Cytogenetics Core at MD Anderson.

Xenografts generated with the D423 (*MTAP*-WT), U87 (*MTAP*-deleted), U87 pCMV MTAP (*MTAP*-rescued); NB1 (*MTAP*-WT), *SK-MEL-5* (*MTAP*-deleted), Gli56 (*MTAP*-deleted) cancer cell lines were generated as detailed below for FFPE sections. For sub-cutaneous xenografts, 5 million cells were injected in the flanks of Nude a thymic mice (bred by M.D. Anderson’s Department of Experimental Radiation Oncology); Tumors were harvested and fixed in 4% phosphate buffered formalin. Dehydration, paraffin embedding and tissue sectioning was performed by M.D. Anderson’s Veterinary Pathology Core. Intracranial glioma cell injections were performed by the M.D. Anderson Intracranial Injection Core at M.D. Anderson (Dr. Fred Lang, Director (Lal et al., 2000)). Intracranial tumors were established by injection of 200,000 cells into the brains of immunocompromised female nude *Foxn1*nu/nu mice ages 4-6 months, which were bred at the Experimental Radiation Oncology Breeding Core in M.D. Anderson. The animals were first bolted and allowed to recover for 2 weeks. Then, glioma cells were injected through the bolt with a Hamilton syringe. Bolting and cell injections were performed by the M.D. Anderson Intracranial Injection Fee-for-Service Core (Lal et al., 2000). All animal procedures were approved by the M.D. Anderson Animal Care and Use Committee. Formalin fixed brains with xenografted glioma tumors were submitted to the Veterinary Pathology Core for embedding and sectioning.

### Metabolomic Profiling of Polar Metabolites

Polar metabolites were profiled using the Beth Israel Deaconess Medical center (BIDMC) platform. Extraction of samples was performed in house as follows for each specific set of samples. Polar metabolites were profiled using the Beth Israel Deaconess Medical center (BIDMC) platform. Extraction of samples was performed in house as follows for each specific set of samples.

#### Targeted Mass Spectrometry

Samples were re-suspended using 20 μL HPLC grade water for mass spectrometry. 5-7 μL were injected and analyzed using a hybrid 5500 QTRAP triple quadrupole mass spectrometer (AB/SCIEX) coupled to a Prominence UFLC HPLC system (Shimadzu) via selected reaction monitoring (SRM) of a total of 262 endogenous water-soluble metabolites for steady-state analyses of samples. Some metabolites were targeted in both positive and negative ion mode for a total of 298 SRM transitions using positive/negative ion polarity switching. ESI voltage was +4950V in positive ion mode and –4500V in negative ion mode. The dwell time was 3 ms per SRM transition and the total cycle time was 1.55 seconds. Approximately 10-14 data points were acquired per detected metabolite. Samples were delivered to the mass spectrometer via hydrophilic interaction chromatography (HILIC) using a 4.6 mm i.d × 10 cm Amide XBridge column (Waters) at 400 μL/min. Gradients were run starting from 85% buffer B (HPLC grade acetonitrile) to 42% B from 0-5 minutes; 42% B to 0% B from 5-16 minutes; 0% B was held from 16-24 minutes; 0% B to 85% B from 24-25 minutes; 85% B was held for 7 minutes to re-equilibrate the column. Buffer A was comprised of 20 mM ammonium hydroxide/20 mM ammonium acetate (pH=9.0) in 95:5 water:acetonitrile. Peak areas from the total ion current for each metabolite SRM transition were integrated using MultiQuant v2.1 software (AB/SCIEX)

### Intracellular metabolites from cells in culture (cell pellet)

Adherent cancer cell lines growing in 10-cm plates at 50-90% confluency were harvested for metabolomic profiling as follows. First, media was removed (and extracted for its own profiling) and the plated cells were washed twice with ice cold 1x PBS (Corning). Cells were then placed in the dry ice and we added 4ml of 80% methanol pre-cooled to −80°C. Then, incubated in the − 80°C freezer for 20 min. Cells were scrapped off the plate while on dry ice, placed in pre-cooled tubes and finally centrifugated at maximum speed, 17,000 × g for 5 minutes at 4°C. The supernatant containing polar metabolites was concentrated by speedvac-ing using an Eppendorf Vacufuge plus (Yuan et al., 2012) and submitted to the BIDMC Mass spec core.

### Secreted metabolites (conditioned media)

1-2ml of conditioned media was centrifuged at max speed for 10 minutes to remove any unblocked cells or cell debris. Subsequently, 4 volumes of −80°C pre-cooled methanol for every 1 volume of media was added to make the final concentration of 80% (vol/vol) methanol solution, then vortexed, and left at −80°C for 6-8 hours to precipitate serum and extract polar metabolites. Ice cold methanol media mix, was centrifugedat maximum speed (17,000 × g) for 10 min at 4°C. Following centrifuging, the supernatant was separated and dried in the speedvac (Eppendorf Vacufuge plus), and was sent to the BIDMC core (Yuan et al., 2012).

### Frozen tumors

Snap liquid nitrogen frozen tumors were kept in the −80°C until extraction and never thawed. Every effort was taken to keep tumors frozen until placed in cold methanol. Frozen tumors were weighed on dry ice without thawing, and placed them in dry ice pre-cooled Fisher Tube #02-681-291 with a Qiagen steel beads. Then, we added 1 ml of 80% at −80°C pre-cooled methanol to each tube. Then, we shacked the tubes with Qiagen Tissue Lyser for 45 s at room temperature at 28 Hz for multiple rounds. We placed the tubes in the dry ice between each round. After samples became homogenous, the final volume of samples were proportioned to 50mg tissue/2ml of 80% methanol. After this step, we kept the samples in −80°C for 24hrs and then vortexed each sample and centrifuged at maximum speed (17,000 × g) for 10 min at 4°C. Then we collect the supernatant and store them in −80°C until speedvacing. For each study, we used Eppendorf Vacufuge Plus to speedvac the same volume of samples. Dried samples were sent to the BIDMC core.

### Determination of MTA by ^1^H NMR in conditioned cell culture media

Conditioned media from glioma cells at 50% confluency was collected and centrifuged for 10 minutes to remove cells and cell debris. Post-centrifugation media was transferred to a new tube, where 4 volumes of methanol were added to precipitate proteins. The sample was then vortexed and centrifuged at 4000*g for 30 minutes and the supernatant was transferred to an Eppendorf tube. The post-centrifuged supernatant was then speedvacced using the Eppendorf Vacufuge Plus and the resulting pellet was resuspended in 750 μL D-DMSO (dimethyl sulfoxide-d6, Alfa Aesar, 99.5% A16893-18). Any insoluble material was removed by centrifugation. The clear D-DMSO solution was then placed in a 5-mm NMR tube for NMR analysis. ^1^H NMR measurements were performed on a Brucker Avance III HD 500 MHz equipped with a cryoprobe broadband observe probe at MD Anderson Cancer Center. The resulting spectrum was obtained using the zg30 pulse sequence with either 100 or 1000 scans and the relaxation delay equal to 1s. Spectrum analysis was performed using Bruker TopSpin 3.1 software. For MTA, the adenosine protons (HMDB0001173), with chemical shifts of 8.15 ppm and 8.35 ppm, were readily detectable in the supernatant of *MTAP*-deleted media extract, but not *MTAP*-intact or *MTAP*-rescued and co-cultured *MTAP*-deleted and *MTAP*-rescued media extract. The most intense peak at of MTA at 2.1 ppm corresponded to the methyl group protons adjacent to the sulfur. However, in the media extract of *MTAP*-deleted cells, this peak is obscured by more abundant metabolites. The peaks for the ribose group were low intensity and not detectable. We validated our observed chemical shifts by spiking sample extracts with pure MTA standard (Sigma Aldrich D5011-100MG).

### Validation of an anti-MTAP rabbit monoclonal antibody for the Immunohistochemistry detection of *MTAP*-homozygously deleted tumors

In order to score homozygous deletions of MTAP by immunohistochemistry in primary tumor FFPE sections, we first validated a monoclonal antibody for this application by demonstrating staining in FFPE slides of xenografted tumors with known MTAP genotype. Xenografts generated with the D423 (*MTAP*-WT), U87 (*MTAP*-deleted), U87 pCMV MTAP (*MTAP*-rescued) NB1 (*MTAP*-WT), *SK-MEL-5* (*MTAP*-deleted), Gli56 (MTAP-deleted) cancer cell lines were fixed in formalin and embedded in paraffin. Immunohistochemistry was performed on coronal sections of mouse brain xenografted with human glioblastoma cells. Tissue was fixed in formaldehyde before being embedded in paraffin. Tissue was sliced to desired thickness and embedded onto slide. Slides were left to incubate at 60°C overnight. For detection of MTAP, sections were subjected antigen retrieval in citrate buffer [1:100 Vector Antigen Unmasking Solution (Citrate-Based) H-3300 250 mL] at [CUISINART HIGH TEMP] for 10 minutes. Sections were covered with a blocking buffer of 2% goat serum [Vector S-1000 Normal Goat Serum 20 mL] in PBS [Quality Biological PBS (10X), pH 7.4 1000 mL) for 1 hour. Sections were then covered with anti-MTAP rabbit monoclonal (Abcam product number: ab126623; clone: EPR6892) in a 1:200 dilution with 2% goat serum in PBS and left to incubate overnight in 4°C. Sections were then washed in PBS and incubated for 30 minutes with [Invitrogen by Thermo Scientific 1X goat anti-rabbit IgG secondaryantibody, poly HRP conjugate]. After washing in PBS and Tween20 [Fisher BioReagents BP337-500], Sections were devloped using either Impact? NOVAred (Vector Labs; yields a red-to brown color for stain) or using EnzMet^tm^ (Nanoprobes #6001-30 mL; yields a black stain). For NOVAred, slides were then counterstained using hematoxylin; for EnzyMet, slides were counterstained with hemoatoxylin and optionally, eosin counterstain. Sections were mounted using Denville Ultra Microscope Cover Glass (Cat# M1100-02) and Thermo Scientific Cytoseal 60 and left to dry overnight. There was an absolute correspondence between the genotype of the xenografts and the MTAP-staining by IHC; with only mouse stromal cells staining positive for MTAP in MTAP-deleted xenografts (**Supplementary Figure S5–S7**). We thus proceeded to utilize this antibody for scoring MTAP homozygous deletions in human primary GBM FFPE sections. IHC was performed as described for the human xenografted tumors. Immunohistochemistry was performed for utilizing citrate antigen retrieval and blocking in 2% goat serum. slides were stained against MTAP (Abcam, rabbit monoclonal ab126623 1:1000, overnight 4°C), and developed with 1x goat anti-rabbit IgG secondary, poly HRP conjugate: thermo scientific, B40962 for 30min at room temp and developed with NOVAred (Vector Labs), or EnzMet (Nanoprobes #6001-30 mL) followed by counterstaining with hematoxylin, or hematoxylin and Eosin.

## ACKNOWLEDGEMENTS

This work was financially supported by the following grants to F.L.M: US National Institutes of Health (NIH) grant 1R21CA226301, the American Cancer Society Research Scholar Award RSG-15-145-01-CDD, the National Comprehensive Cancer Network – Young Investigator Award YIA170032 and the Andrew Sabin Family Foundation Fellows Award. The University of Texas MD Anderson Cancer Center/Glioblastoma Moon Shot, The CABI/GE In-Kind Research Grant (MI2), the Brockman Medical Research Foundation and the SPORE in Brain Cancer (2P50CA127001) funds, also supported the work. We thank Lisa Norberg and Kristin Alfaro-Munoz for GBM FFPE slide preparation and Edward Chang for slide scanning.

## AUTHOR CONTRIBUTIONS

Y.B. performed metabolomic sample preparation, NMR and data analysis; J.J.A. and N.B.S. performed western blots; J.J.A., K.A, T.T., and A.H.P. performed immunohistochemistry, Y.L. performed molecular biology and cell culture. A.H.P, T.T., D.K.G., and E.S.B. generated figures and analyzed data. J.T.H. analyzed IHC slides. J.D.G., J.T.H., A.C and R.V. shared genomic data and primary tumor samples; J.M.A performed metabolomics analysis; V.C.Y., Y.B. and F.L.M. conceived the study and wrote the paper.

## Supplemental Information

**Supplementary Figure S1:**
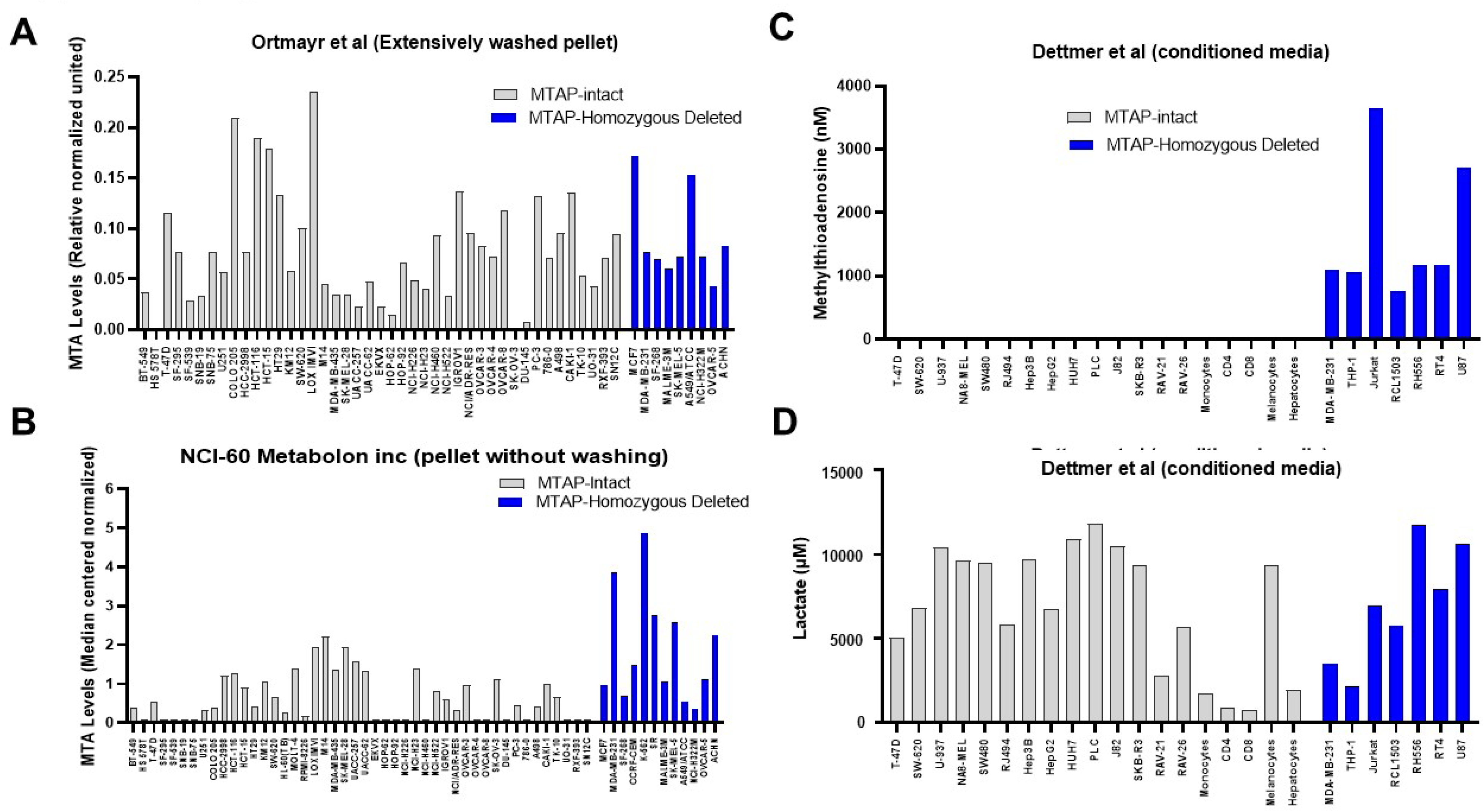
Measurements of MTA levels of *MTAP*-intact versus deleted cancer cell line vary depending on how well intracellular versus secreted metabolites are separated during sample preparation. Methylthioadenosine (MTA) levels from three different studies (Ortmayr et al., 2019; Dettmer et al., 2013; Su et al., 2011) are compared. Briefly, MTA levels from the supplementary data of those studies are re-plotted, with each bar representing the levels of a single cell line. Most of these cell lines are part of the NCl-60 panel (cell line name in x-axis, *MTAP*-deleted cell lines in blue). The study by Ortmayr et al (Panel A) specified that the cell pellets were washed extensively, whilst that from NCl-60 (performed by Metabolon Inc, Panel B) did not. Even though both studies were conducted with the same set of cell lines and under the same culturing conditions, the NCl-60 Metabolon Inc data show that the MTAP-deleted cell lines have a -3-fold higher MTA levels(0.62± 0.09 vs 1.87 ±0.4, P=0.01, t-test), whilst the Ortmayr data show no difference (0.077 ± 0.008 vs 0.089 ± 0.014, n.s.). Panel C: The study by Dettmer et al used conditioned media, reflecting MTA levels that are i.e. exclusively extracellular. The levels of MTA are >100-fold higher in *MTAP*-deleted vs *MTAP*-intact cell lines(7.9 ± 1.7 (n=19) vs 1663 ± 408 (n=7), P<0.007 t-test). Panel D: As a control, the lactate levels in the conditioned media from the Dettmer et al study are shown. No difference is seen between the *MTAP* deleted and intact groups. The MTA levels are given as average ± SEM., the t-test is with unpaired, two tails, unequal variance.

**Supplementary Figure S2:**
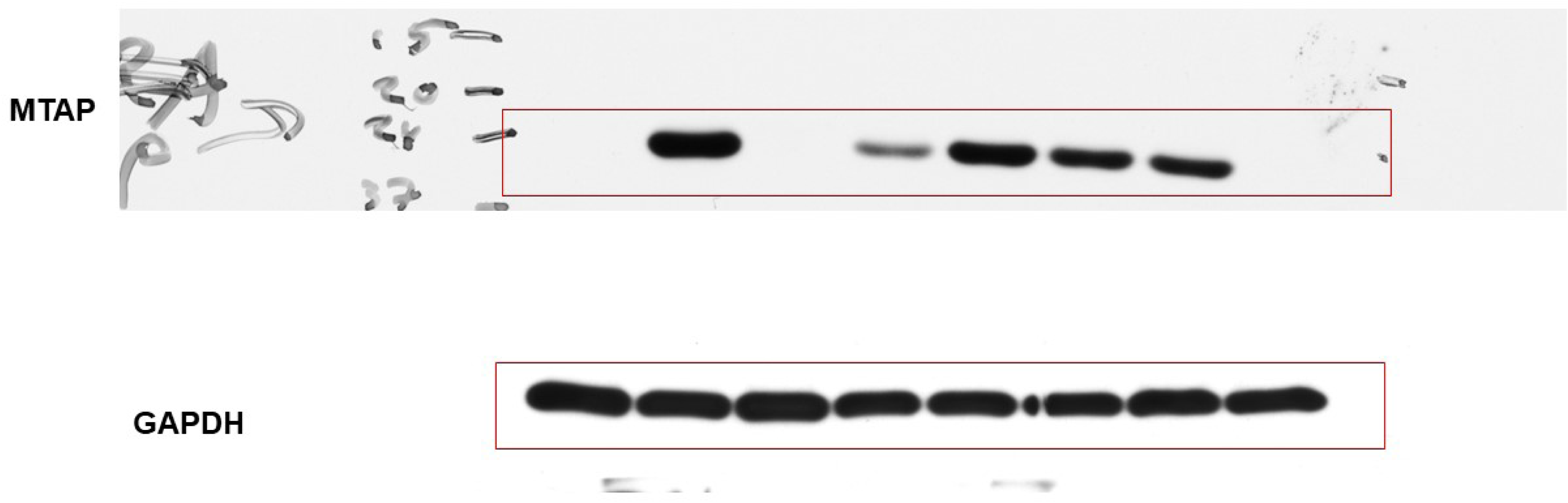
Western blot raw scans for Figure 2.

**Supplementary Figure S3:**
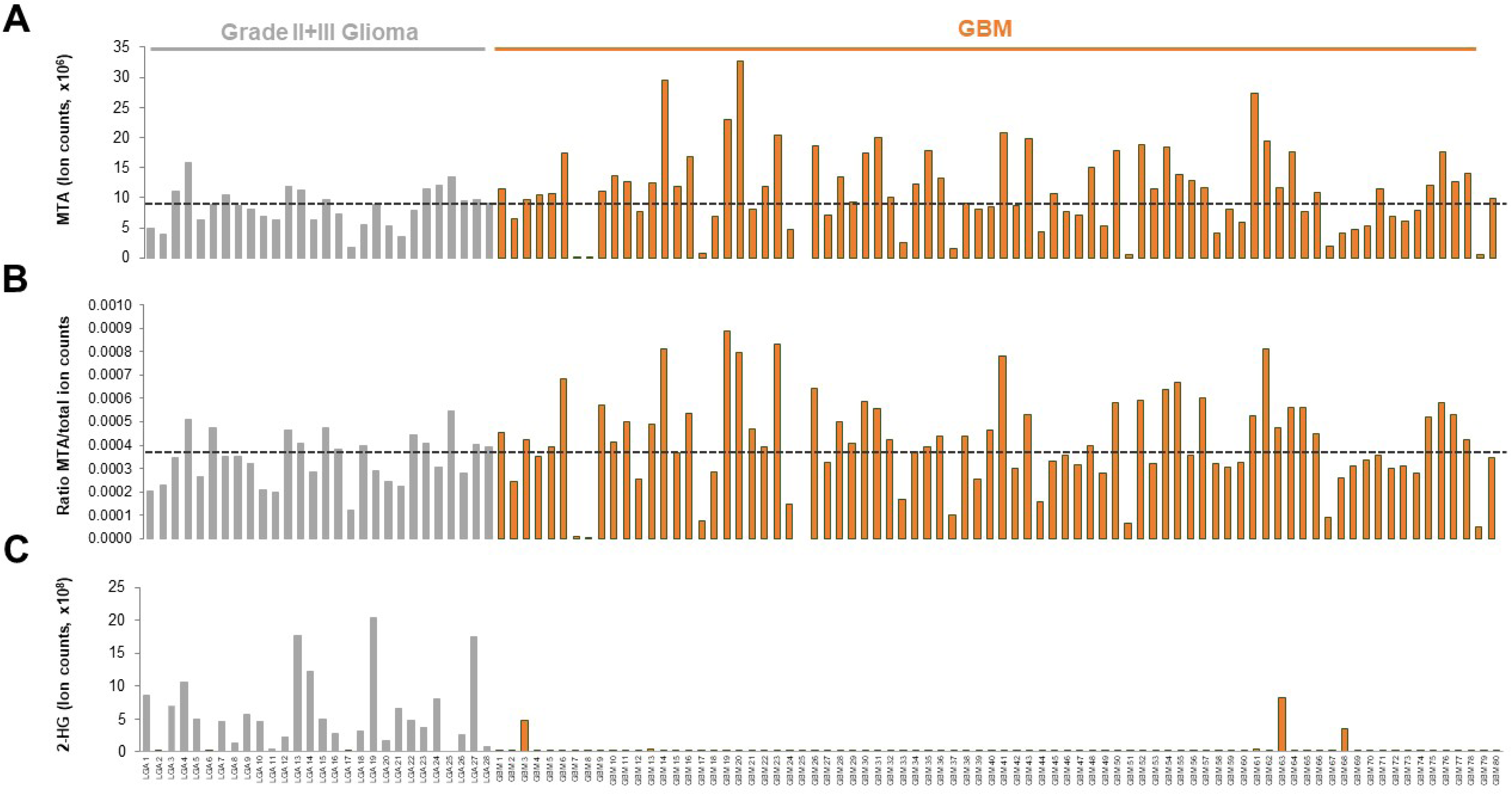
Metabolomic profiling studies of primary human tumors show minimal differences in MTA between Lower Grade Glioma and Grade IV GBM despite dramatically higher incidence of *MTAP*-deletions in GBM. **Panel A:** MTA levels from a Metabolon Inc profile study (*Neuro-Oncology*, Volume 21, Issue 3, March 2019, Pages 337–347); MTA levels were replotted from Supplementary data of this study. **Panel B:** MTA levels from the same study expressed as a ratio of total ion count to account for differential loading. **Panel C:** Levels of 2-HG in the same dataset, highlighting what a massive increase in a specific metabolite by a specific genetic event looks like, and how it contrasts with the marginal increases in MTA in *MTAP*-deleted tumors.

**Supplementary Figure S4:**
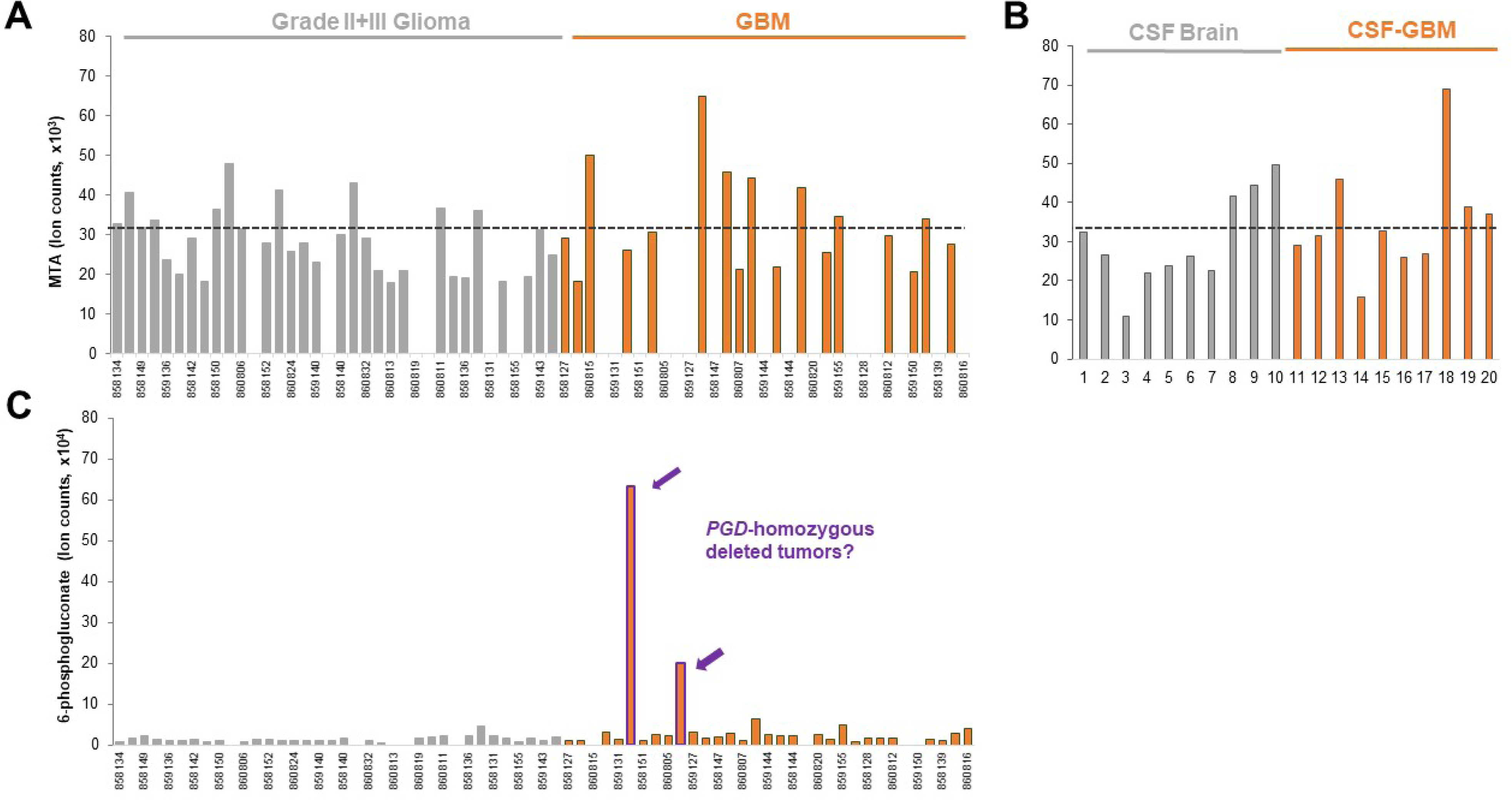
Minimal elevation of MTA in primary GBM tumors or CSF. **Panel A:** MTA levels in a series of primary low-grade gliomas (grey) and GBM (orange) from a global metabolome profile using the Metabolon Inc platform, plotted form Supplementary data from ref Chinnaiyan *et al*. Cancer Res. 2012. The median levels of MTA in the LGG gliomas, an approximation for *MTAP* WT tumors, is plotted in a dashed line. **Panel B.** 6-phosphogluconate (6-PG) levels from the same study. Tumors with exceptionally high levels of 6-PG are very likely to have 1p36 homozygous deletions that includes *PGD*. **Panel C:** MTA levels in cerebrospinal fluid (CSF) collect from GBM versus normal brain, from a metabolomics profile using the BIDMC platform (Locasale *et al*. Mol Cell Proteomics 2012). The median levels of MTA in normal brain CSF is shown by a grey line. While the *MTAP*-deletion status of individual tumors in that study is not given, ~40% of GBM tumors have *MTAP*-homozygous deletions, so it is near certain that *MTAP*-deleted cases are part of this collection. Assuming that the cases with highest levels of MTA are in fact those with *MTAP* deletions, this means that *MTAP*-deleted GBM tumors have at best, 2-fold higher levels of MTA in CSF as compared to normal brain. The same reasoning can be applied to the data in **Panel A and Figure S3A**; at best, there is a 2-fold increase in MTA in *MTAP*-deleted compared to the median of *MTAP-WT* GBM.

**Supplementary Figure S5:**
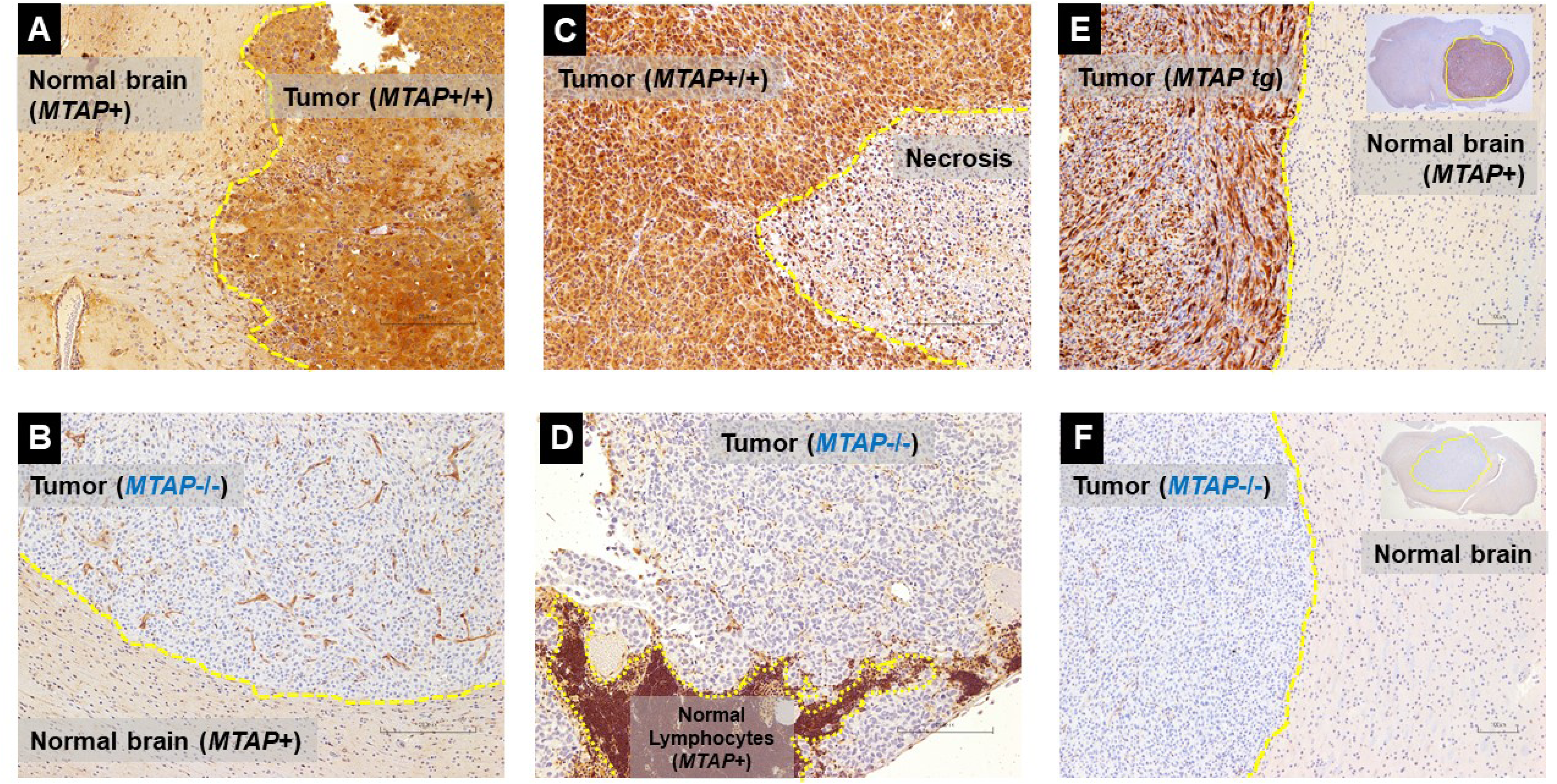
Validation of an MTAP rabbit monoclonal antibody for the detection of *MTAP*-deleted tumors by IHC on FFPE sections. Xenografted tumors (S.C.: A, B; intracranial: C,D, E,F) differencing in MTAP-deletion status (**A**, D423, MTAP intact, **B,** U87, *MTAP*-homozygous deleted, **C,** NB1, *MTAP*-intact, SK-MEL-5, *MTAP*-homozygous deleted, **E**, U87 pCMV MTAP; *MTAP*-rescued, **F**, U87 *MTAP*-homozygous deleted) were grown in immunocompromised mice and FFPE sections generated. IHC was performed with rabbit monoclonal anti-MTAP (ab126623; EPR6892) and slides developed by NOVA red (red-brown staining indicating MTAP presence) and counterstained by hematoxylin (blue, nuclei). Tumor boundaries are shown in yellow. Note the clear correspondence between *MTAP* genomic status and staining intensity in tumors, with complete absence of staining in MTAP-deleted tumors. The MTAP antibody yields strong staining pretty much in all cells except those with *MTAP*-deletions and regions of necrosis. This fully validates that this genuinely detects MTAP protein by IHC in FFPE sections.

**Supplementary Figure S6:**
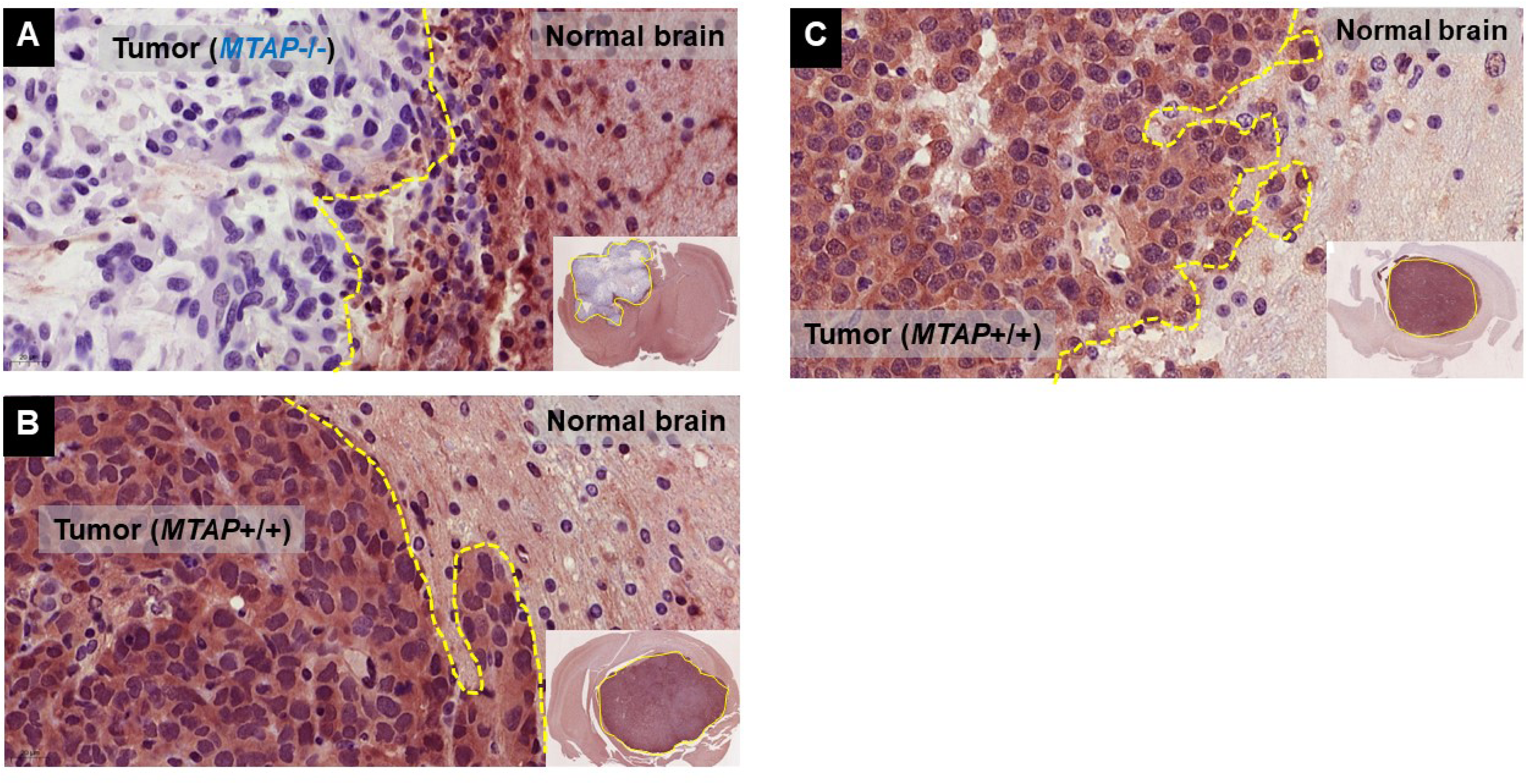
IHC Staining with the MTAP monoclonal antibody remains specific even at longer exposures. FFPE sections of xenografted tumors generated from glioma cell lines differing in *MTAP* deletion status were stained with anti-MTAP rabbit monoclonal (ab126623) and developed with NOVAred. **Panel A:** Gli56 (*MTAP*−/−; deleted); **B:** NB1, C:0423 *(*MTAP*+I+*; intact); The exposure of the developer was increased compared to experiments in Fig S5, in order to determine whether non-specific background staining would start to occur in *MTAP*-deleted tumors; no evidence of this is present (Panel A), fully validating the use of this antibody to evaluate *MTAP*-deletion status in human FFPE GBM sections.

**Supplementary Figure S7:**
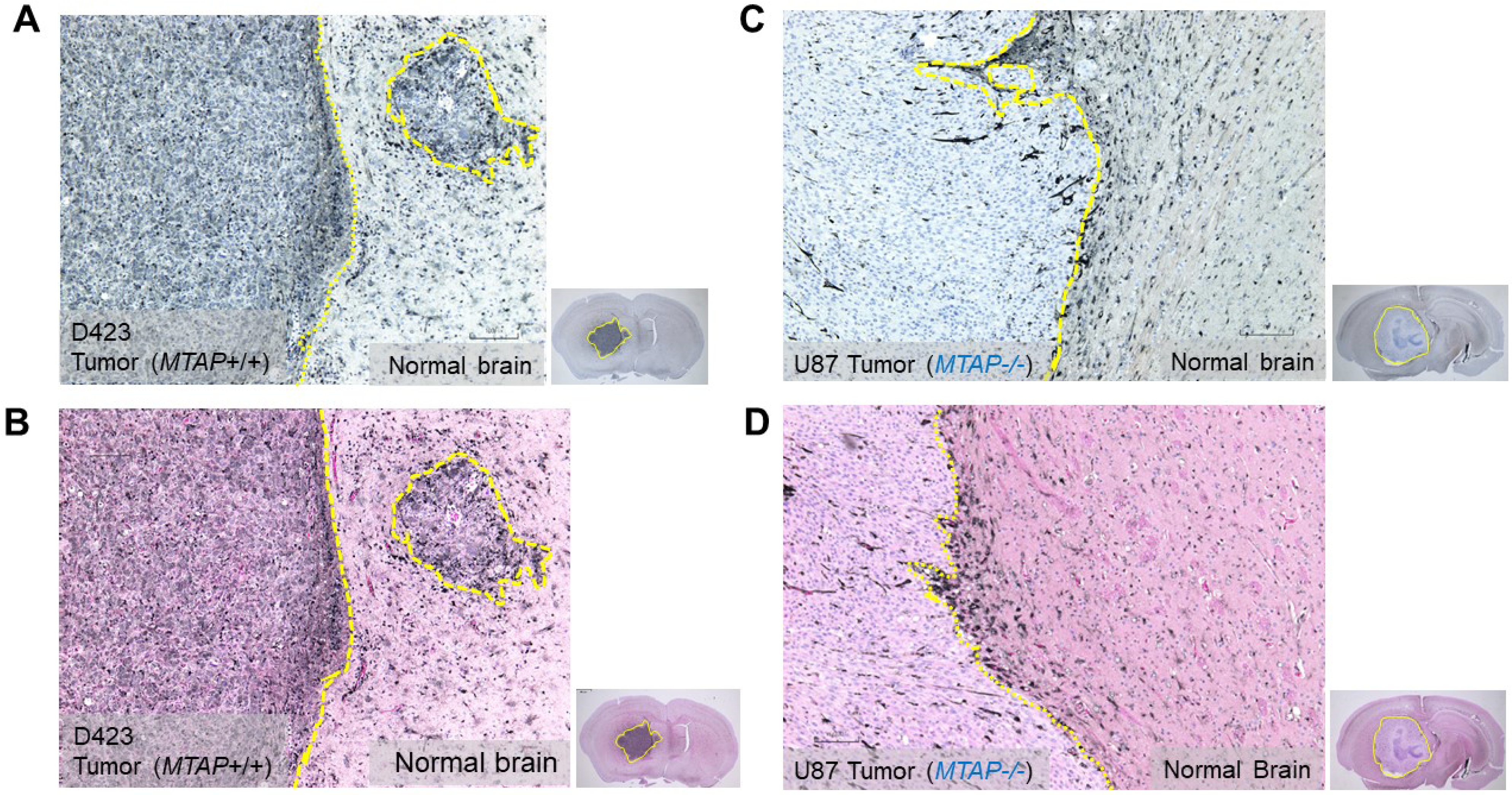
MTAP IHC developed by EnzMet with or without Eosin counterstain yields increased histological resolution. **A, B:** lntracranial xenografts generated with 0423 (*MTAP-WT*) glioma cells, **C, D:** with the U87 (*MTAP*-deleted) glioma cell lines. IHC staining with anti-MTAP (ab126623) was performed as in Fig S2 and Fig S3, except that instead of NovaRed, antibodies were developed using the EnzMet silver developer Areas of immunopositivity (MTAP presence) are black rather than Red/brown in the case of NovaRED. Unlike NOVARED or DAB, EnzMet staining is not washed out by ethanol, allowing Eosin staining for higher level of histological detail to be discernable. **A, C**; stained with anti-MTAP and developed by EnzyMet with Hematoxylin counterstain and **B, D** with additional Eosin counterstain. The advantage of EnzyMet are lower background, and higher resolution, as well as tolerance to Eosin counterstain. We have used this developer for the FFPE primary human GBM studies (Figure 4).

**Supplementary Figure S8:**
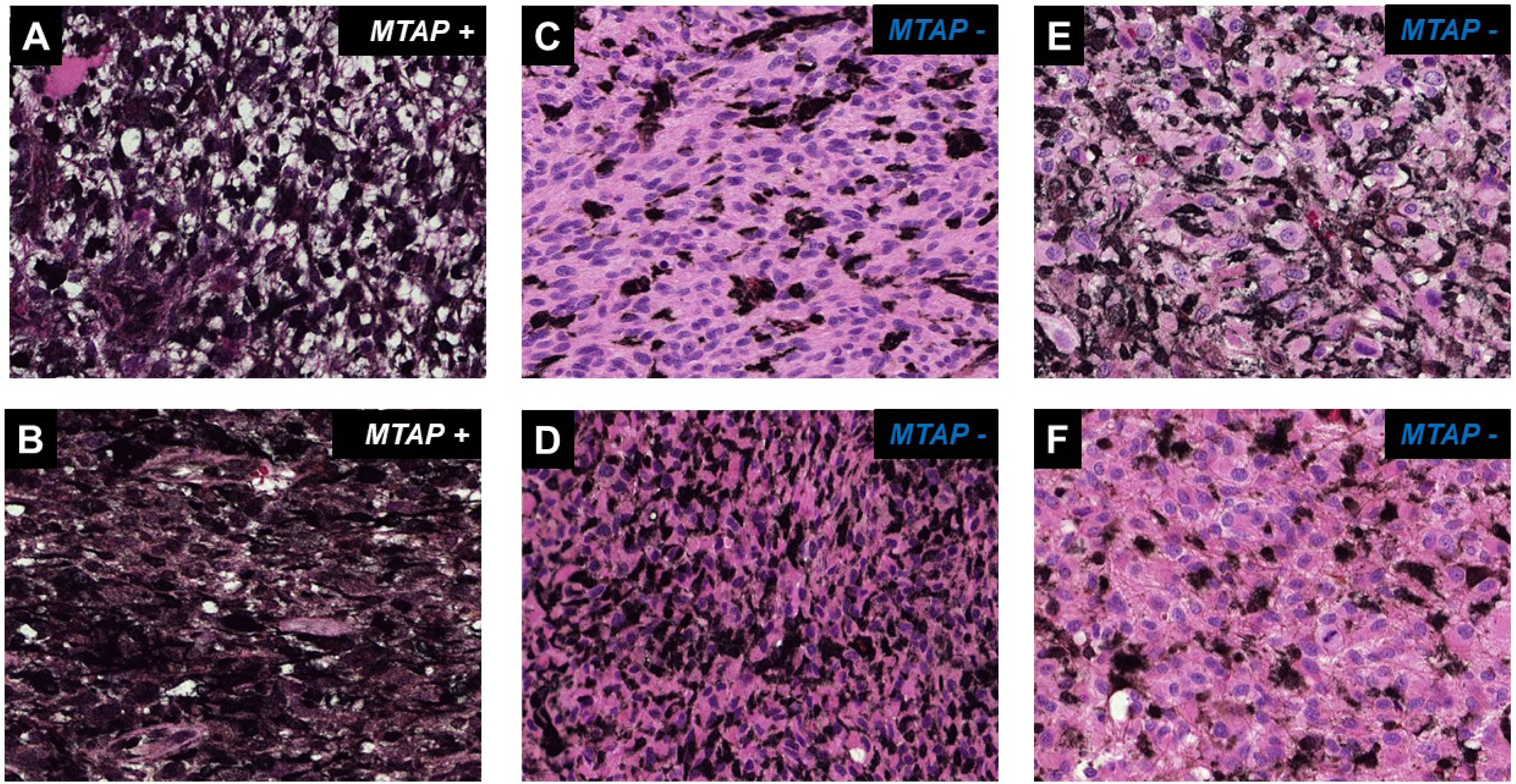
Representative GBM tumors stained with anti-MTAP antibody differing in stromal content. Experiments were performed as in Figure 4 and images acquired using a 40X objective. Representative cases of *MTAP* positive and *MTAP* negative GBM tumors that differ in the percent of stromal content are shown. **Panel A:** Case 882732, *MTAP* positive; **Panel B:** Case 902391, *MTAP* positive; **Panel C:** Case 958196 *MTAP* negative; **Panel D:** Case 9100168: *MTAP* negative, this tumor is an example of extreme stromal content approaching 50 % cellularity. **Panel E:** Case 880244, *MTAP* negative; **Panel F:** Case 999128, *MTAP* negative; this tumor is an example of minimal stromal content.

**Supplementary Figure S9:**
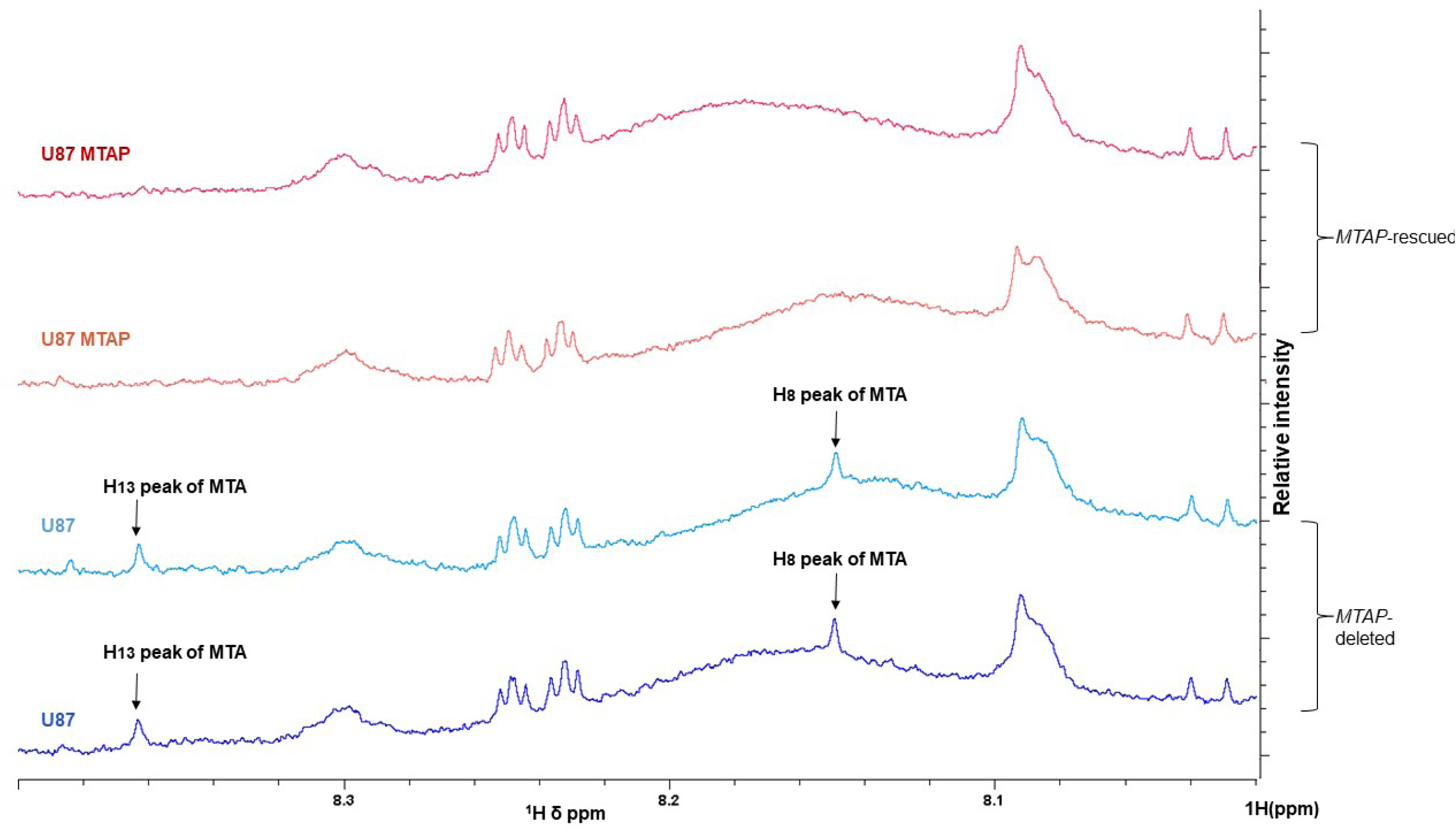
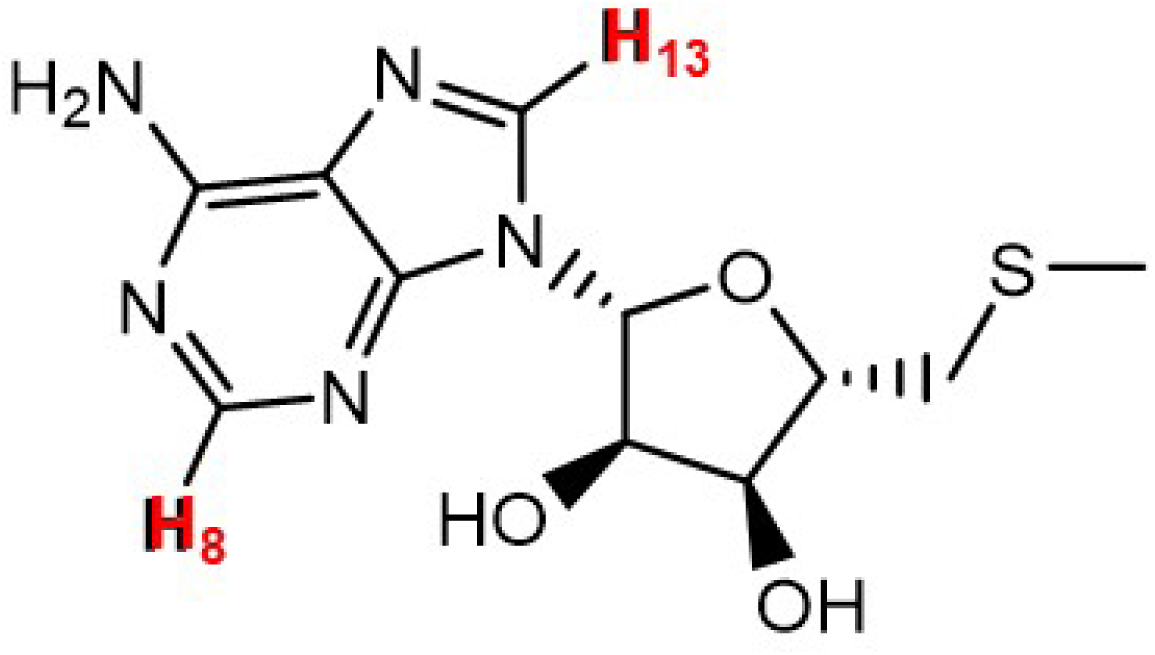
^1^H NMR detection of MTA in polar extracts of conditioned media from *MTAP*-deleted but not *MTAP*-rescued glioma cells. Conditioned media from *MTAP*-homozygously deleted U87 and *MTAP*-rescued U87 glioma cells was collected after 3 days of growth (−80% confluency). 2 ml of media was extracted with 80% methanol as described in methods and the extract was dissolved in 500 µL of 06 deuterated-DMSO. Samples were analyzed in a 500 MHz Brucker NMR (1H, NS=100 transients). The spectral region from 8ppm-8.5ppm is shown, and spectra were normalized to an invariant metabolite (8.1 ppm). The two peaks corresponding to H13 and H7 hydrogen atoms of MTA (HMDB0001173) appear in the U87 (*MTAP*-deleted) media extract, while they are absent in U87 MTAP (*MTAP*-rescued) media extract. Identity of these peaks to MTA was confirmed by spiking with a pure standard. Other ^1^H peaks of MTA in the 4-2 ppm chemical shift were obscured by more abundant metabolites in media. Two independent biological replicates are shown for each cell line.

**Supplementary Figure S10:**
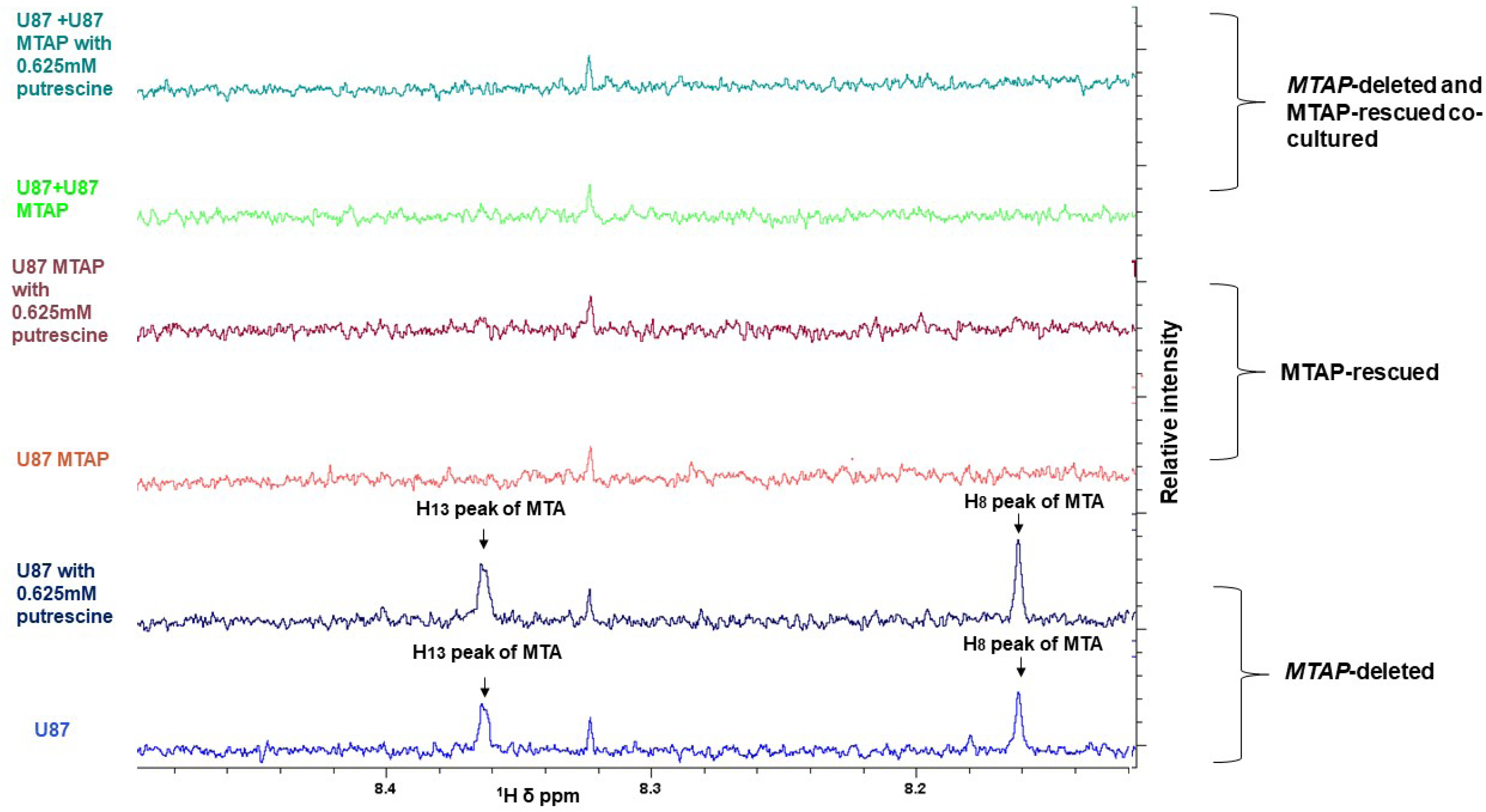
Co-culture of *MTAP*-deleted and *MTAP*-rescued glioma cells abrogates accumulation of MTA in conditioned media. Equal numbers of U87 and U87 *MTAP*-rescued cells were co-cultured for 3 days with or without treatment with 0.625 mM putrescine (to stimulate MTA format ion, see Figure 1b; Kamatani and Carson, Cancer Res. 1980) after which media was extracted with ethyl acetate (see methods). NMR spectra of U87 and U87 MTAP and U87 + U87 MTAP media extracts were acquired by a 500 MHz NMR (NS=1000) treated with or without putrescine the immediate precursor of methylthioadenosine (MTA, see Figure 1b) and dissolved in 0.5 ml of Deuterated 06-DMSO (1000 transients). The spectral region from 8 ppm-8.5 ppm is shown, and spectra are normalized to the internal metabolites. The two peaks of H13 and H8 of MTA appear in the U87 and U87 treated with putrescine media extract, while they are absent in U87 MTAP and U87+U87 MTAP media extract. Other peaks of MTA did not show up in the U87 media extract because either they have low intensity, or they are obscured with other abundant metabolites in media. This figure proves that in U87 cells MTA is secreted out into the media. Also, it shows putrescine increases the amount of MTA in media.

