## Supplemental Info for "Methylthioadenosine is Not Dramatically Elevated in *MTAP*-Homozygous Deleted Primary Glioblastomas"

**Supplementary Figure S1:** Measurements of MTA levels of *MTAP*-intact versus deleted cancer cell line vary depending on how well intracellular versus secreted metabolites are separated during sample preparation

**Supplementary Figure S2:** Western blot raw scans for Figure 2

**Supplementary Figure S3:** Metabolomic profiling studies of primary human tumors show minimal differences in MTA between Lower Grade Glioma and Grade IV GBM despite dramatically higher incidence of *MTAP*-deletions in GBM

**Supplementary Figure S4**: Minimal elevation of MTA in primary GBM tumors or CSF

**Supplementary Figure S5:** Validation of an MTAP rabbit monoclonal antibody for the detection of *MTAP*-deleted tumors by IHC on FFPE sections

**Supplementary Figure S6:** IHC staining with the MTAP monoclonal antibody remains specific even at longer exposures.

**Supplementary Figure S7:** MTAP IHC developed by EnzMet with or without Eosin counterstain yields increased histological resolution

**Supplementary Figure S8:** Representative GBM tumors stained with anti-MTAP antibody differing in stromal content

**Supplementary Figure S9:** $^{1}$H NMR detection of MTA in polar extracts of conditioned media from *MTAP*-deleted but not *MTAP*-rescued glioma cells

**Supplementary Figure S10:** Co-culture of *MTAP*-deleted and *MTAP*-rescued glioma cells abrogates accumulation of MTA in conditioned media

### Supplementary Figure S1

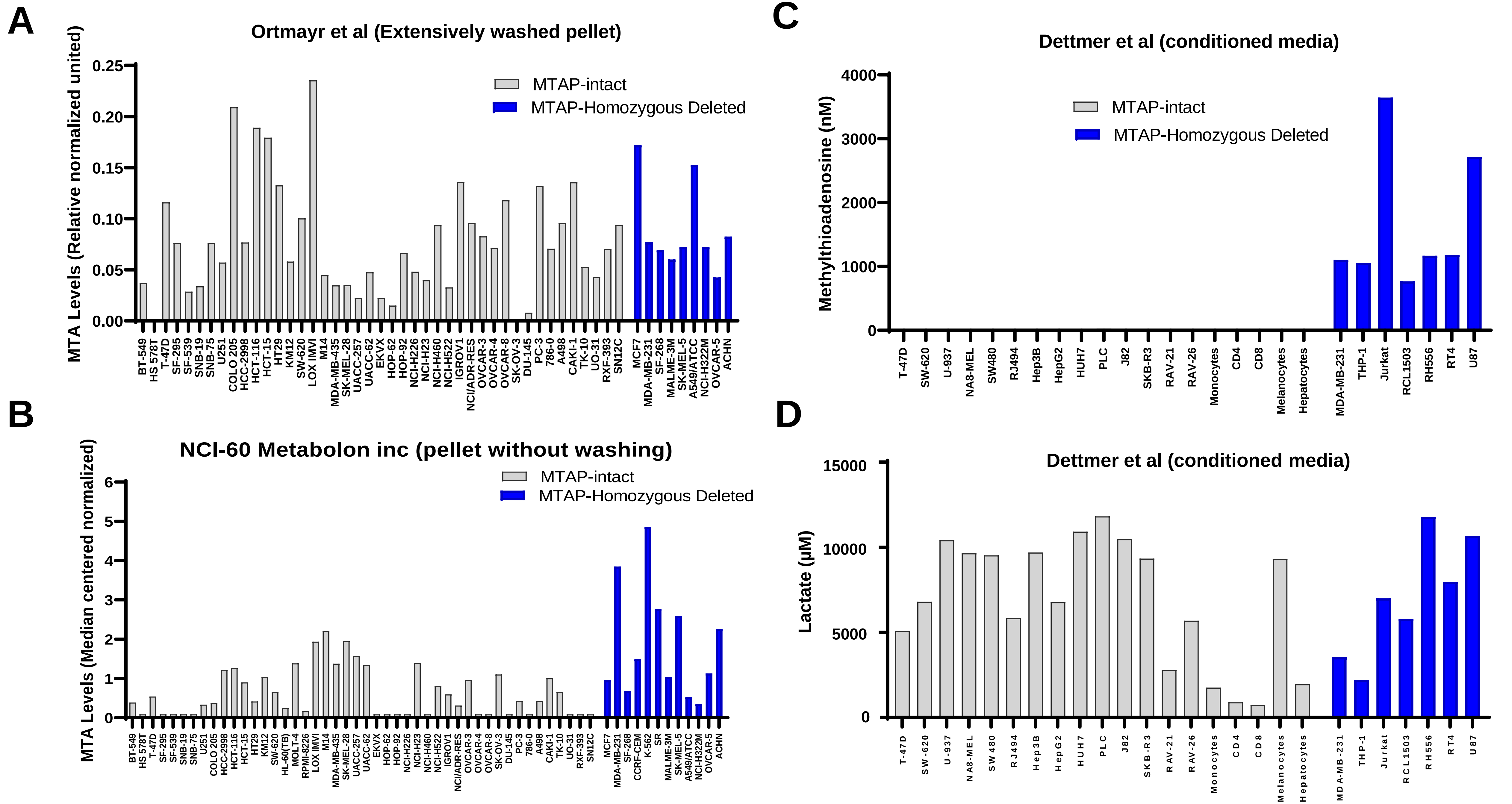

**Supplementary Figure S1: Measurements of MTA levels of *MTAP*-intact versus deleted cancer cell line vary depending on how well intracellular versus secreted metabolites are separated during sample preparation.** Methylthioadenosine (MTA) levels from three different studies (Ortmayr et al., 2019; Dettmer et al., 2013; Su et al., 2011) are compared. Briefly, MTA levels from the supplementary data of those studies are re-plotted, with each bar representing the levels of a single cell line. Most of these cell lines are part of the NCI-60 panel (cell line name in x-axis, *MTAP*-deleted cell lines in blue). The study by Ortmayr et al (Panel A) specified that the cell pellets were washed extensively, whilst that from NCI-60 (performed by Metabolon Inc, Panel B) did not. Even though both studies were conducted with the same set of cell lines and under the same culturing conditions, the NCI-60 Metabolon Inc data show that the *MTAP*-deleted cell lines have a ~3-fold higher MTA levels(0.62± 0.09 vs 1.87 ±0.4, P=0.01, t-test) , whilst the Ortmayr data show no difference (0.077 ± 0.008 vs 0.089 ± 0.014, n.s.). Panel C: The study by Dettmer et al used conditioned media, reflecting MTA levels that are i.e. exclusively extracellular. The levels of MTA are >100-fold higher in *MTAP*-deleted vs *MTAP*-intact cell lines(7.9 ± 1.7 (n=19) vs 1663 ± 408 (n=7), P<0.007 t-test ). Panel D: As a control, the lactate levels in the conditioned media from the Dettmer et al study are shown. No difference is seen between the *MTAP* deleted and intact groups. The MTA levels are given as average ± SEM., the t-test is with unpaired, two tails, unequal variance.

#### Supplementary Figure S2

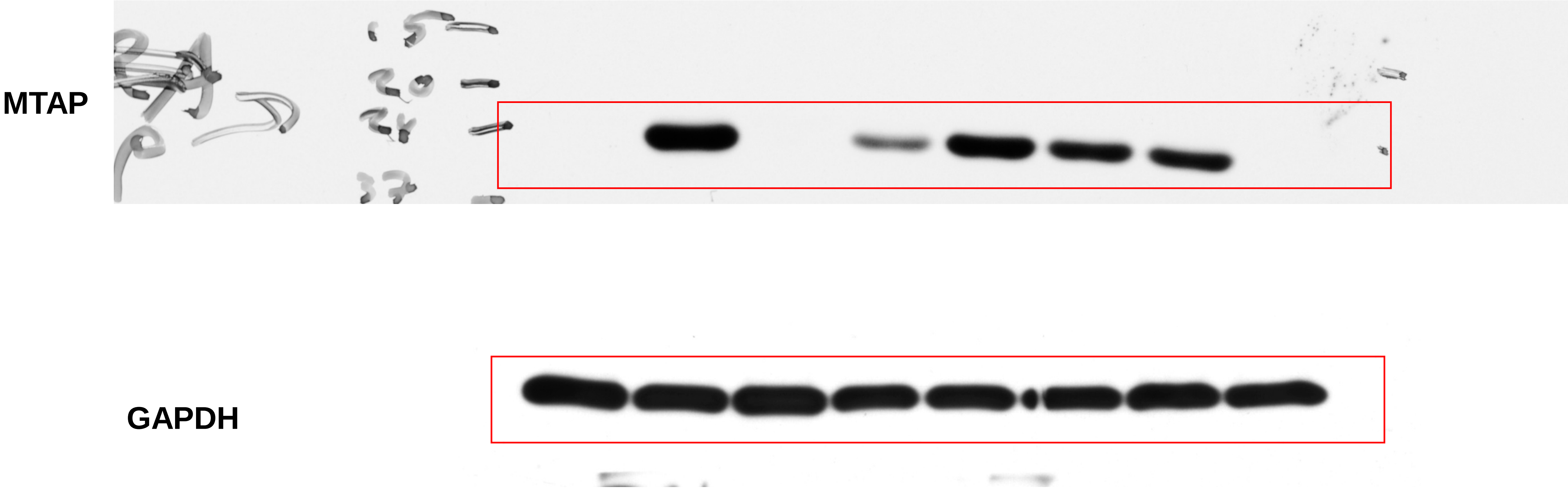


**Supplementary Figure S2: Western blot raw scans for Figure 2**

#### Supplementary Figure S3

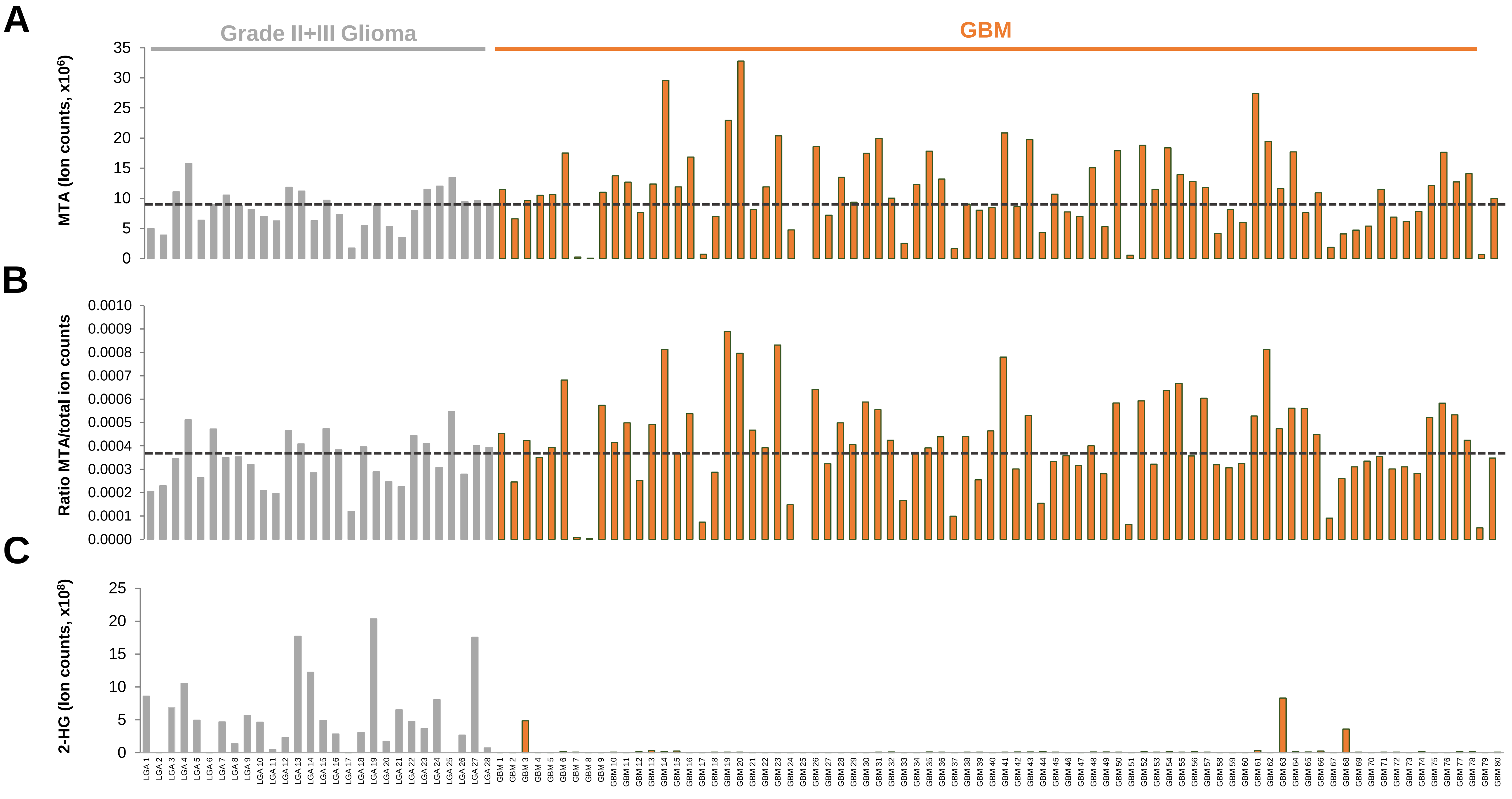

**Supplementary Figure S3: Metabolomic profiling studies of primary human tumors show minimal differences in MTA between Lower Grade Glioma and Grade IV GBM despite dramatically higher incidence of *MTAP*-deletions in GBM.**

**Panel A:** MTA levels from a Metabolon Inc profile study (*Neuro-Oncology*, Volume 21, Issue 3, March 2019, Pages 337–347); MTA levels were replotted from Supplementary data of this study. **Panel B**: MTA levels from the same study expressed as a ratio of total ion count to account for differential loading. **Panel C**: Levels of 2-HG in the same dataset, highlighting what a massive increase in a specific metabolite by a specific genetic event looks like, and how it contrasts with the marginal increases in MTA in *MTAP*-deleted tumors.

### Supplementary Figure S4

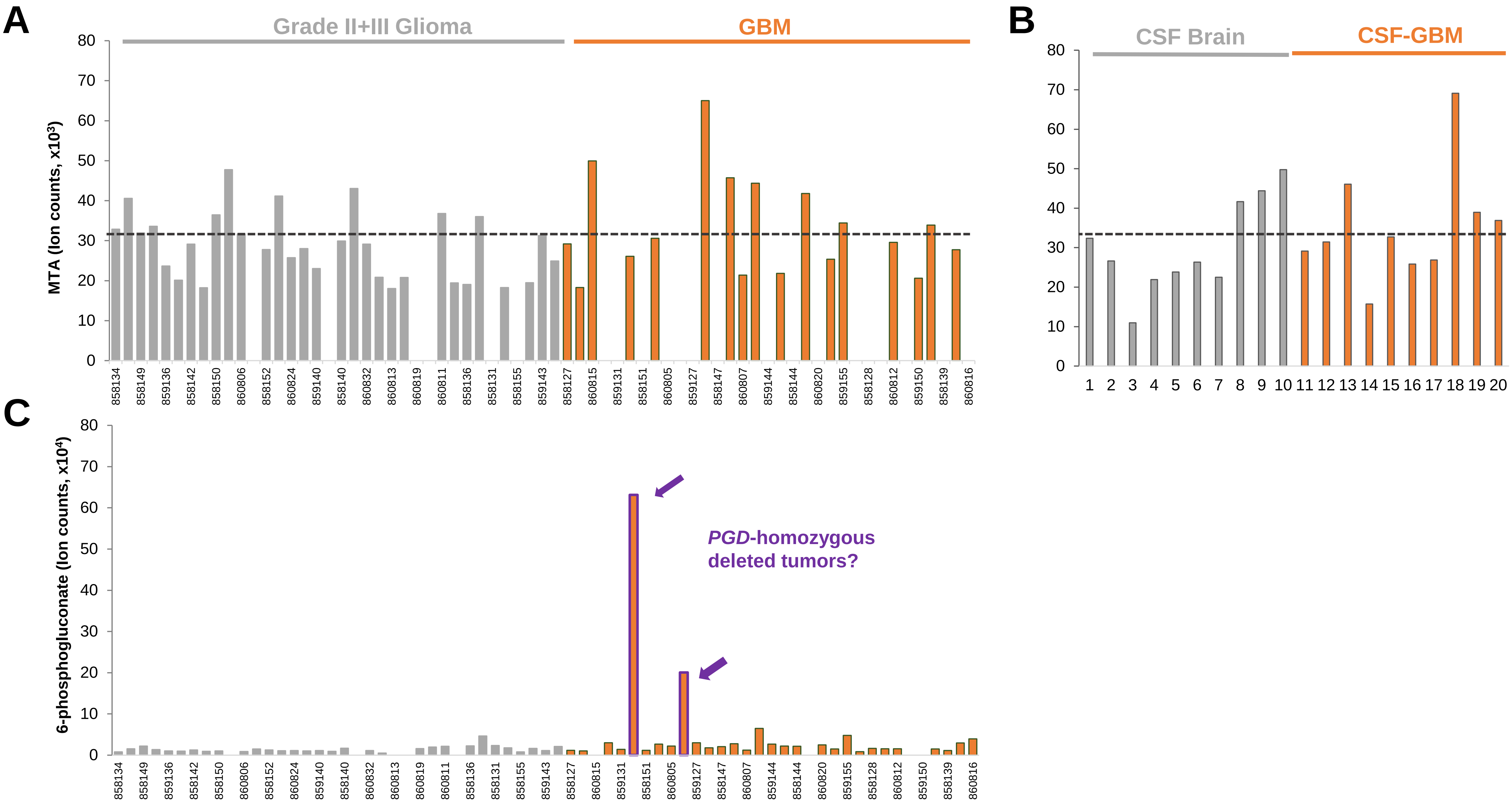

**Supplementary Figure S4: Minimal elevation of MTA in primary GBM tumors or CSF**

**Panel A:** MTA levels in a series of primary low-grade gliomas (grey) and GBM (orange) from a global metabolome profile using the Metabolon Inc platform, plotted form Supplementary data from ref Chinnaiyan *et al.* Cancer Res. 2012. The median levels of MTA in the LGG gliomas, an approximation for *MTAP* WT tumors, is plotted in a dashed line. **Panel B**. 6-phosphogluconate (6-PG) levels from the same study. Tumors with exceptionally high levels of 6-PG are very likely to have 1p36 homozygous deletions that includes *PGD*. **Panel C:** MTA levels in cerebrospinal fluid (CSF) collect from GBM versus normal brain, from a metabolomics profile using the BIDMC platform (Locasale *et al.* Mol Cell Proteomics 2012). The median levels of MTA in normal brain CSF is shown by a grey line.

While the *MTAP*-deletion status of individual tumors in that study is not given, ~40% of GBM tumors have *MTAP*-homozygous deletions, so it is near certain that *MTAP*-deleted cases are part of this collection. Assuming that the cases with highest levels of MTA are in fact those with *MTAP* deletions, this means that *MTAP*-deleted GBM tumors have at best, 2-fold higher levels of MTA in CSF as compared to normal brain. The same reasoning can be applied to the data in **Panel A and Figure S3A**; at best, there is a 2-fold increase in MTA in *MTAP*-deleted compared to the median of *MTAP*-WT GBM.

### Supplementary Figure S5

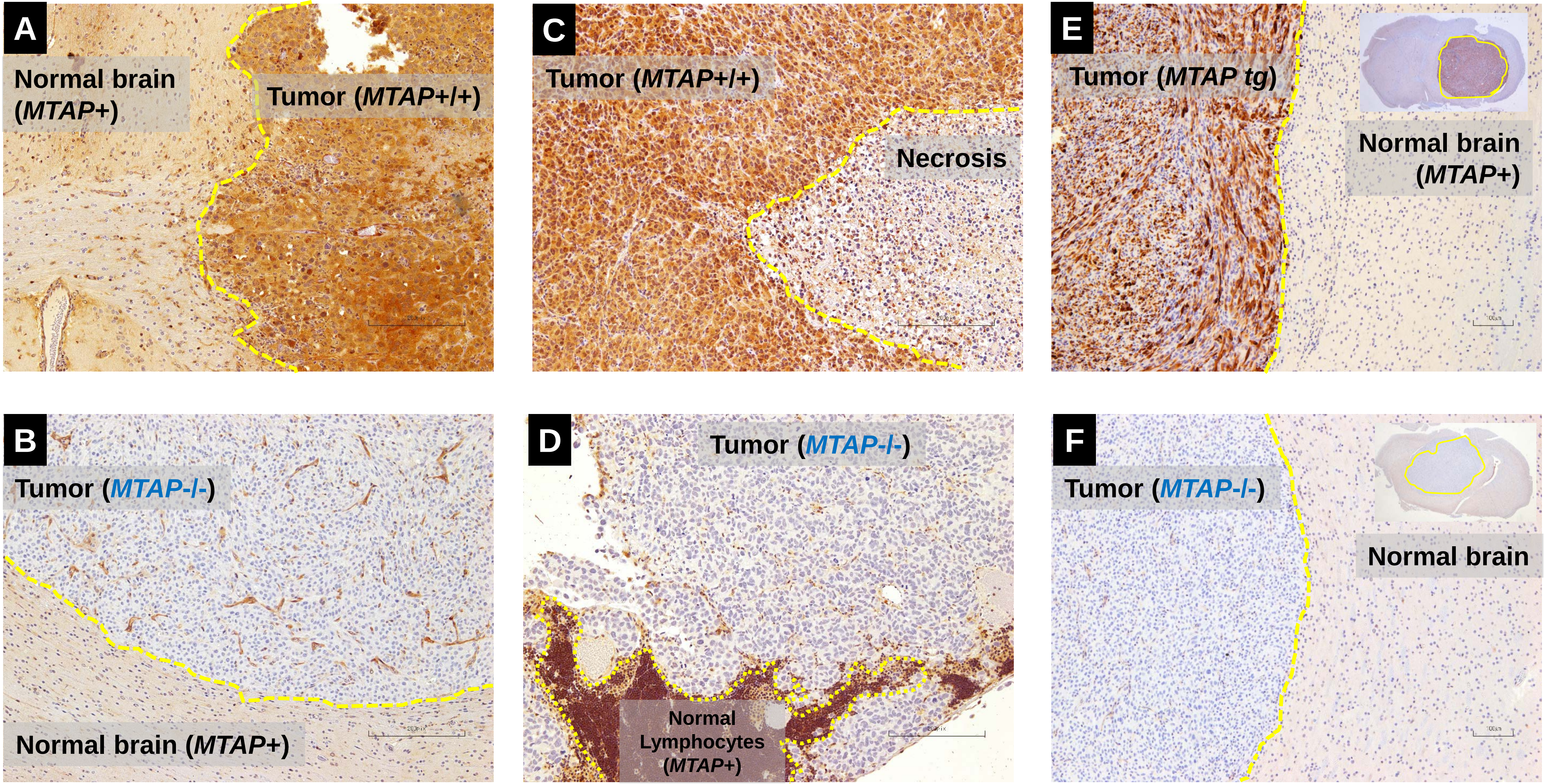

**Supplementary Figure S5: Validation of an MTAP rabbit monoclonal antibody for the detection of *MTAP*-deleted tumors by IHC on FFPE sections** Xenografted tumors (S.C.: A, B; intracranial: C,D, E,F) differencing in *MTAP*-deletion status (**A**, D423, MTAP intact, **B,** U87, *MTAP*-homozygous deleted, **C**, NB1, *MTAP*-intact, **D**, SK-MEL-5, *MTAP*-homozygous deleted, **E**, U87 pCMV MTAP; *MTAP*-rescued, **F,** U87 *MTAP*-homozygous deleted) were grown in immunocompromised mice and FFPE sections generated. IHC was performed with rabbit monoclonal anti-MTAP (ab126623; EPR6892) and slides developed by NOVA red (red-brown staining indicating MTAP presence) and counterstained by hematoxylin (blue, nuclei). Tumor boundaries are shown in yellow. Note the clear correspondence between *MTAP* genomic status and staining intensity in tumors, with complete absence of staining in *MTAP*-deleted tumors. The MTAP antibody yields strong staining pretty much in all cells except those with *MTAP*-deletions and regions of necrosis. This fully validates that this genuinely detects MTAP protein by IHC in FFPE sections.

#### Supplementary Figure S6

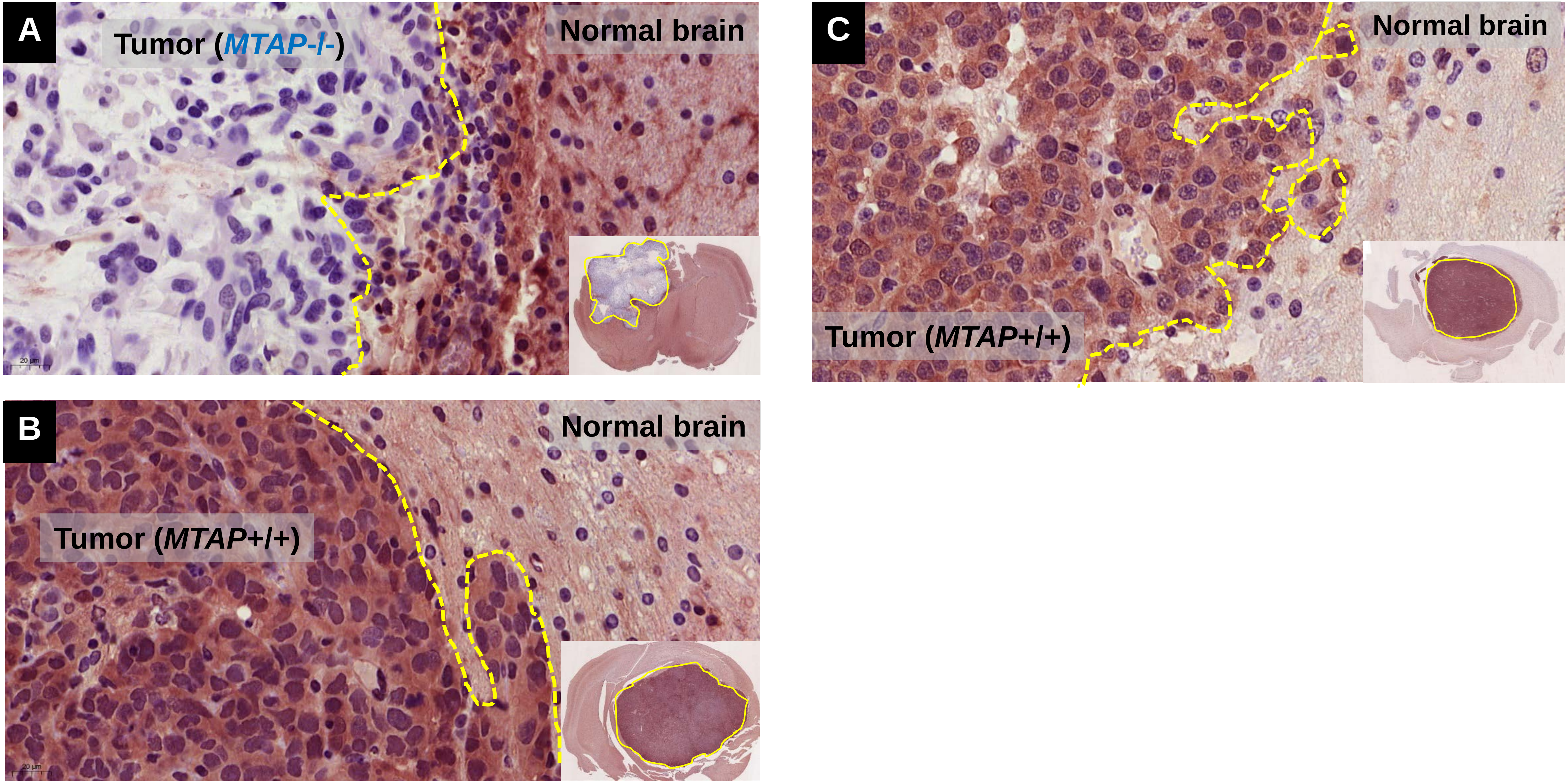

**Supplementary Figure S6: IHC Staining with the MTAP monoclonal antibody remains specific even at longer exposures.** FFPE sections of xenografted tumors generated from glioma cell lines differing in *MTAP* deletion status were stained with anti-MTAP rabbit monoclonal (ab126623) and developed with NOVAred. **Panel A**: Gli56 (*MTAP*-/-; deleted); **B:** NB1, **C:**D423 (*MTAP*+/+; intact); The exposure of the developer was increased compared to experiments in Fig S5, in order to determine whether non-specific background staining would start to occur in *MTAP*-deleted tumors; no evidence of this is present (Panel A), fully validating the use of this antibody to evaluate *MTAP*-deletion status in human FFPE GBM sections.

**Supplementary Figure S7**

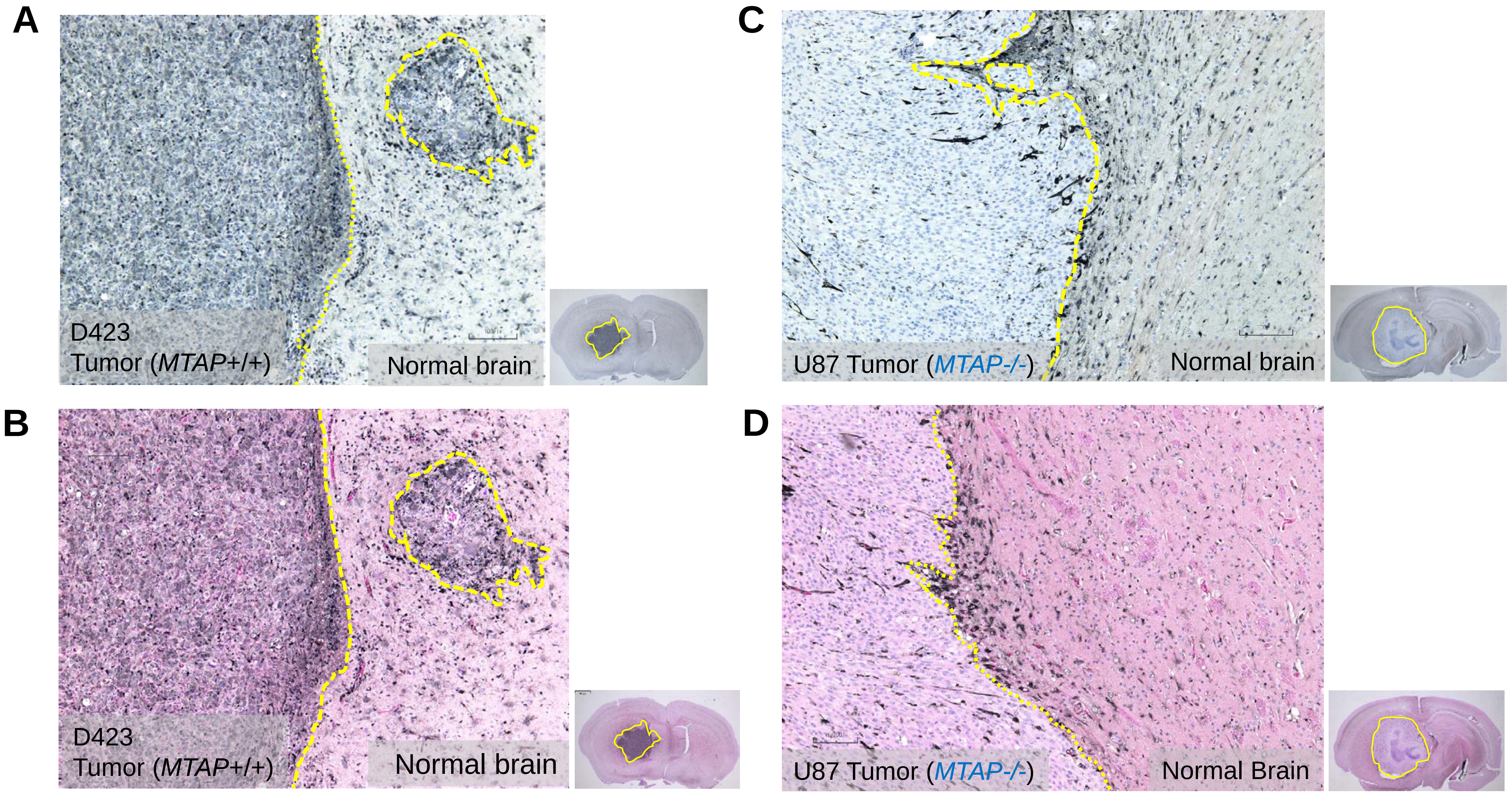

**Supplementary Figure S7: MTAP IHC developed by EnzMet with or without Eosin counterstain yields increased histological resolution**

**A, B**: Intracranial xenografts generated with D423 (*MTAP*-WT) glioma cells, **C, D**: with the U87 (*MTAP*-deleted) glioma cell lines. IHC staining with anti-MTAP (ab126623**)** was performed as in Fig S2 and Fig S3, except that instead of NovaRed, antibodies were developed using the EnzMet silver developer. Areas of immunopositivity (MTAP presence) are black rather than Red/brown in the case of NovaRED. Unlike NOVARED or DAB, EnzMet staining is not washed out by ethanol, allowing Eosin staining for higher level of histological detail to be discernable. **A, C**; stained with anti-MTAP and developed by EnzyMet with Hematoxylin counterstain and **B, D** with additional Eosin counterstain. The advantage of EnzyMet are lower background, and higher resolution, as well as tolerance to Eosin counterstain. We have used this developer for the FFPE primary human GBM studies (Figure 4).

**Supplementary Figure S8**

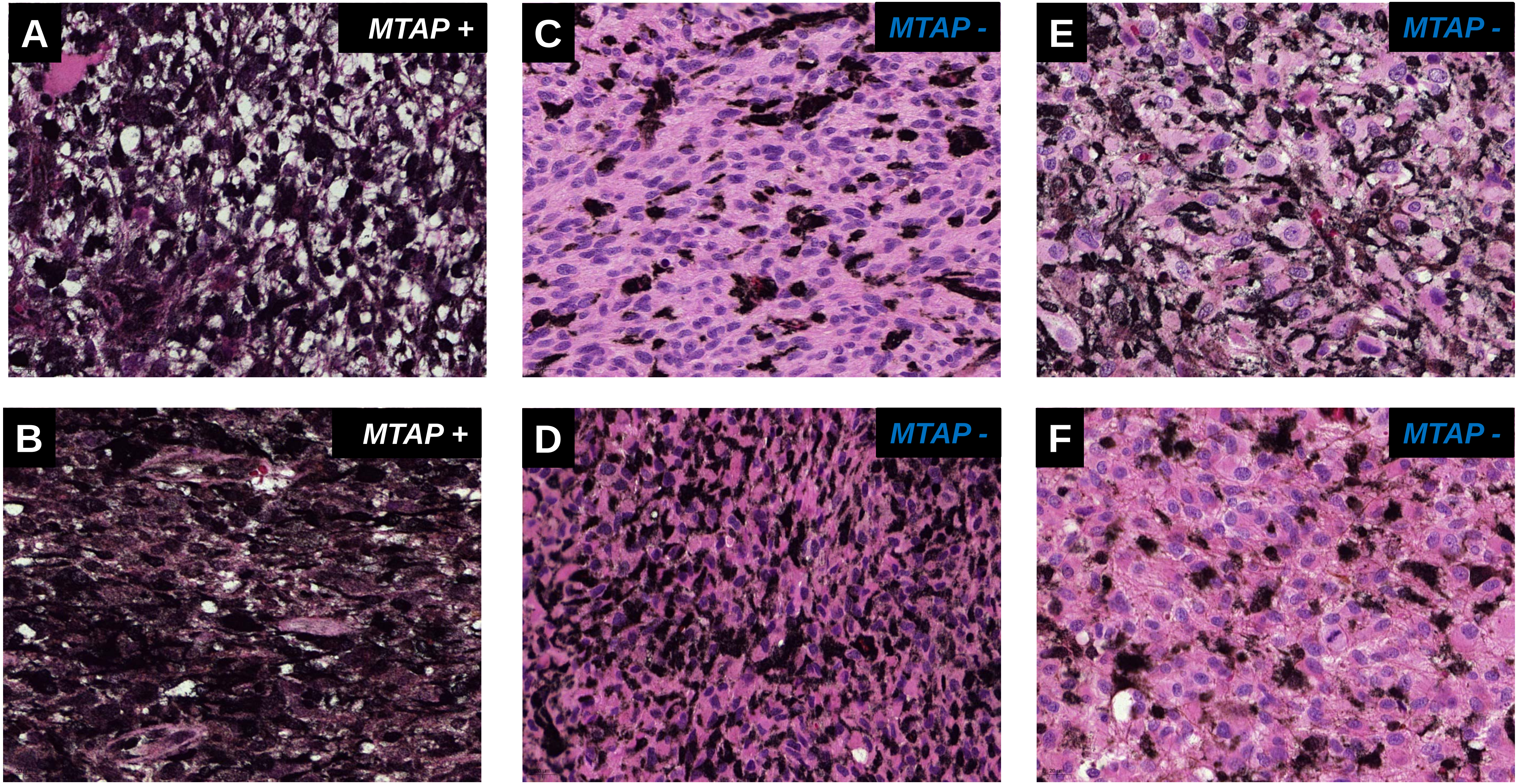

**Supplementary Figure S8: Representative GBM tumors stained with anti-MTAP antibody differing in stromal content.** Experiments were performed as in Figure 4 and images acquired using a 40X objective. Representative cases of *MTAP* positive and *MTAP* negative GBM tumors that differ in the percent of stromal content are shown. **Panel A:** Case 882732, *MTAP* positive; **Panel B**: Case 902391, *MTAP* positive; **Panel C**: Case 958196 *MTAP* negative; **Panel D**: Case 9100168: *MTAP* negative, this tumor is an example of extreme stromal content approaching 50 % cellularity. **Panel E**: Case 880244, *MTAP* negative; **Panel F**: Case 999128, *MTAP* negative; this tumor is an example of minimal stromal content.

#### Supplementary Figure S9

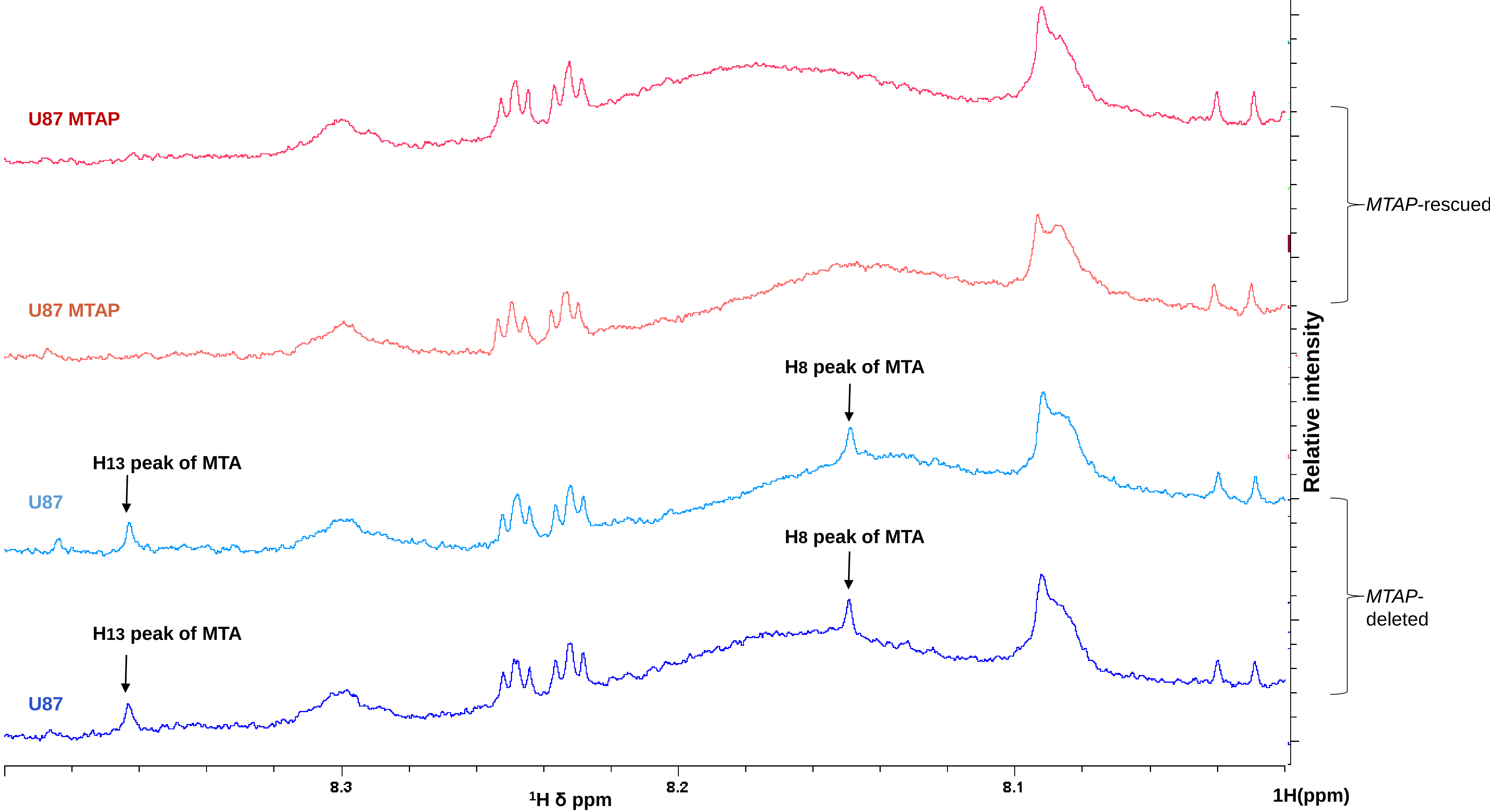

**Supplementary Figure S9: $^1$H NMR detection of MTA in polar extracts of conditioned media from *MTAP*-deleted but not *MTAP*-rescued glioma cells** Conditioned media from *MTAP*-homozygously deleted U87 and *MTAP*-rescued U87 glioma cells was collected after 3 days of growth (~80% confluency). 2 mL of media was extracted with 80% methanol as described in methods and the extract was dissolved in 500 µL of D6 deuterated-DMSO. Samples were analyzed in a 500 MHz Brucker NMR ($^1$H, NS=100 transients). The spectral region from 8ppm-8.5ppm is shown, and spectra were normalized to an invariant metabolite (8.1 ppm). The two peaks corresponding to H13 and H7 hydrogen atoms of MTA (HMDB0001173) appear in the U87 (*MTAP*-deleted) media extract, while they are absent in U87 MTAP (*MTAP*-rescued) media extract. Identity of these peaks to MTA was confirmed by spiking with a pure standard. Other $^1$H peaks of MTA in the 4-2 ppm chemical shift were obscured by more abundant metabolites in media. Two independent biological replicates are shown for each cell line.

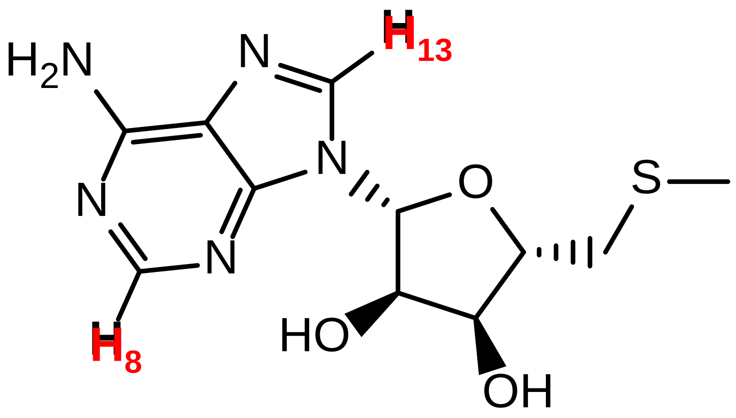

### Supplementary Figure S10

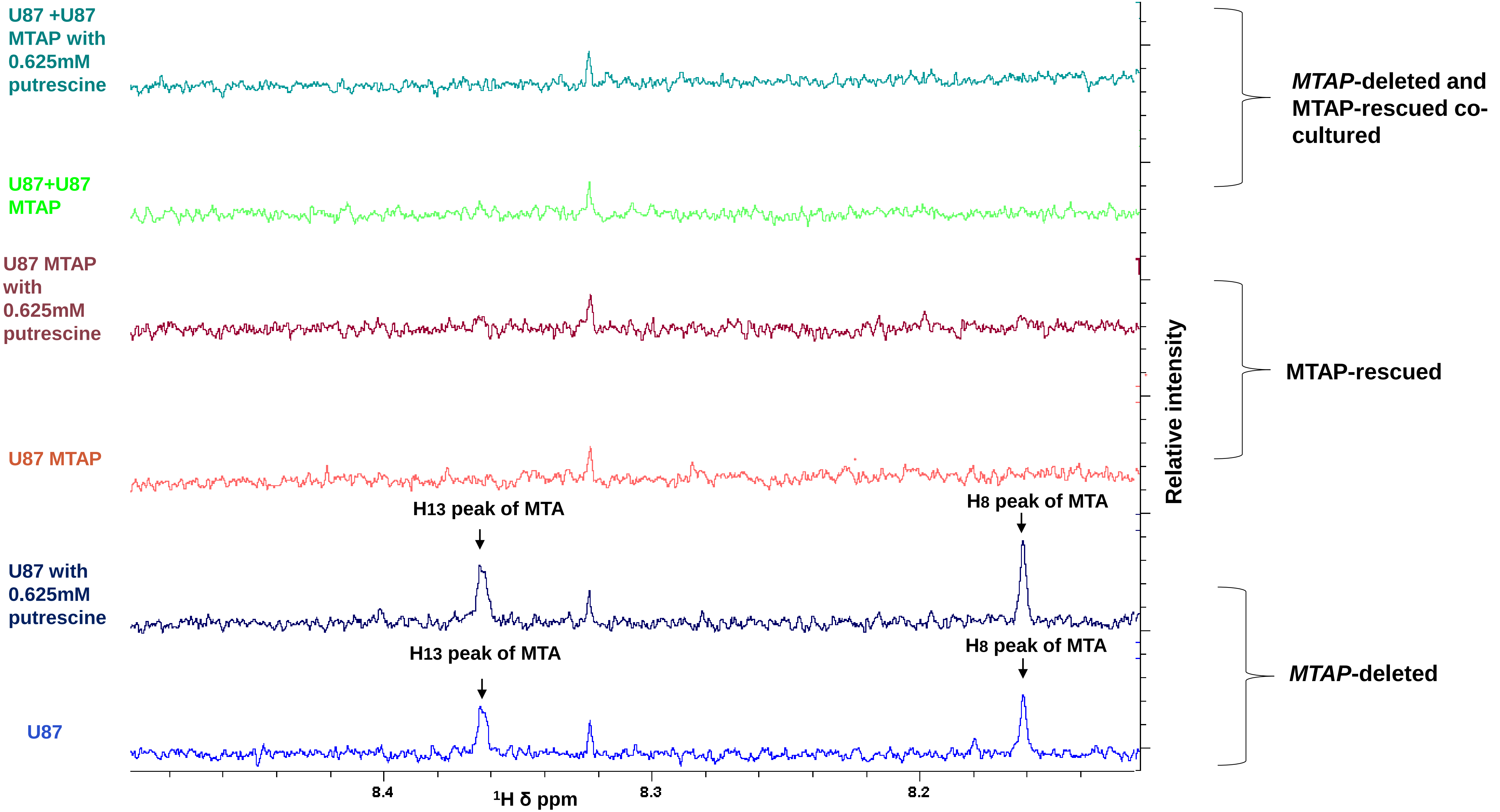

**Supplementary Figure S10: Co-culture of *MTAP*-deleted and *MTAP*-rescued glioma cells abrogates accumulation of MTA in conditioned media** Equal numbers of U87 and U87 *MTAP*-rescued cells were co-cultured for 3 days with or without treatment with 0.625 mM putrescine (to stimulate MTA formation, see Figure 1b; Kamatani and Carson, Cancer Res. 1980) after which media was extracted with ethyl acetate (see methods). NMR spectra of U87 and U87 MTAP and U87 + U87 MTAP media extracts were acquired by a 500 MHz NMR (NS=1000) treated with or without putrescine the immediate precursor of methylthioadenosine (MTA, see Figure 1b) and dissolved in 0.5 ml of Deuterated D6-DMSO (1000 transients). The spectral region from 8 ppm-8.5 ppm is shown, and spectra are normalized to the internal metabolites. The two peaks of H13 and H8 of MTA appear in the U87 and U87 treated with putrescine media extract, while they are absent in U87 MTAP and U87+U87MTAP media extract. Other peaks of MTA did not show up in the U87 media extract because either they have low intensity, or they are obscured with other abundant metabolites in media. This figure proves that in U87 cells MTA is secreted out into the media. Also, it shows putrescine increases the amount of MTA in media.
